# Select autosomal dominant DFNA11 deafness mutations activate Myo7A in epithelial cells

**DOI:** 10.1101/2024.09.17.613491

**Authors:** Prashun Acharya, Garima Thapa, Xiayi Liao, Samaneh Matoo, Maura J. Graves, Sarah Y. Atallah, Ashna K. Tipirneni, Tram Nguyen, Niki. M. Chhabra, Jaden Maschack, Mackenzie R. Herod, Favour A. Ohaezu, Alder Robison, Ashwini Mudaliyar, Jasvinder Bharaj, Nicole Roeser, Katherine Holmes, Vishwaas Nayak, Rayah Alsayed, Benjamin J. Perrin, Scott W. Crawley

**Affiliations:** Department of Biological Sciences, University of Toledo, Toledo, Ohio, USA; Department of Biology, Indiana University, Indianapolis, Indiana, USA

**Keywords:** Myosin, light chains, actin, microvilli, stereocilia, deafness, brush border, epithelia

## Abstract

Myosin-7A (Myo7A) is a motor protein crucial for the organization and function of stereocilia, specialized actin-rich protrusions on the surface of inner ear hair cells that mediate hearing. Mutations in Myo7A cause several forms of genetic hearing loss, including autosomal dominant DFNA11 deafness. Despite its importance, the structural elements of Myo7A that control its motor activity within cells are not well understood. In this study, we used cultured kidney epithelial cells to screen for mutations that activate the motor-dependent targeting of Myo7A to the tips of apical microvilli on these cells. Our findings reveal that Myo7A is regulated by specific IQ motifs within its lever arm, and that this regulation can function at least partially independent of its tail sequence. Importantly, we demonstrate that many of the DFNA11 deafness mutations reported in patients activate Myo7A targeting, providing a potential explanation for the autosomal dominant genetics of this form of deafness.

## INTRODUCTION

Myosin-7A (Myo7A) is a motor protein that belongs to the MyTH4-FERM subfamily of myosins that are typically associated with specialized actin-based protrusions that extend from cells. Myo7A exhibits a relatively restricted tissue expression, found in the inner ear, retina, testis, and kidney (Hasson *et al*., 1995). With the exception of the inner ear, the precise role of Myo7A in most of these tissues remains to be elucidated. In the inner ear, Myo7A is essential for the proper function of stereocilia, large actin-based protrusions on the surface of hair cells that mediate hearing (Self *et al*., 1999; Calabro *et al*., 2019). During development, stereocilia are organized into rows of graded height to form a mechanosensory array known as a hair bundle (Fig 1A). Assembly of a hair bundle is guided by the formation of extracellular linkages that join neighboring stereocilia together, controlling their organization (Muller, 2008). These linkages become more spatially restricted as the hair bundle matures, with a prominent linkage remaining that connects the tips of shorter stereocilia to the side of their taller neighbor (Fig. 1B). This linkage, known as a tip-link, gates a mechanoelectrical transduction (MET) channel complex in response to sound to mediate hearing. A single tip-link filament is composed of two protocadherins that interact via a robust heterophilic adhesion bond (Kazmierczak *et al*., 2007). The upper end of a tip-link filament is cadherin-23 (CDH23), while the lower end is protocadherin-15 (PCDH15). PCDH15 interacts directly with several putative components of the MET channel complex at stereocilia tips (Xiong *et al*., 2012; Maeda *et al*., 2014; Zhao *et al*., 2014; Beurg *et al*., 2015). CDH23 interacts with a cytoplasmic complex composed of two scaffold molecules, USH1C (also known as Harmonin) and USH1G (also known as Sans), as well as Myo7A (Fig. 1B) (Adato *et al*., 2005; El-Amraoui and Petit, 2005). Together, Myo7A and its binding partners form a motor-active complex that “pulls” on the tip-link to maintain a resting tension (Li *et al*., 2020). Importantly, genetic defects in the *MYO7A* gene can result in a number of sensory disorders in humans. Point mutations and in-frame amino acid deletions have been mapped to cause both autosomal recessive (DFNB2) and autosomal dominant (DFNA11) forms of non-syndromic deafness (Friedman *et al*., 2020). Furthermore, mutations, truncations and splice site deletions in the *MYO7A* gene can also result in autosomal recessive Usher Syndrome Type 1B (USH1B), a form of congenital deafness with adolescent-onset progressive blindness (Friedman *et al*., 2020). While autosomal recessive DFNB2 and USH1B mutations are thought to either impair or abolish the activity of Myo7A in cells, how autosomal dominant DFNA11 deafness mutations exert their pathology is unknown.

**Figure 1.**
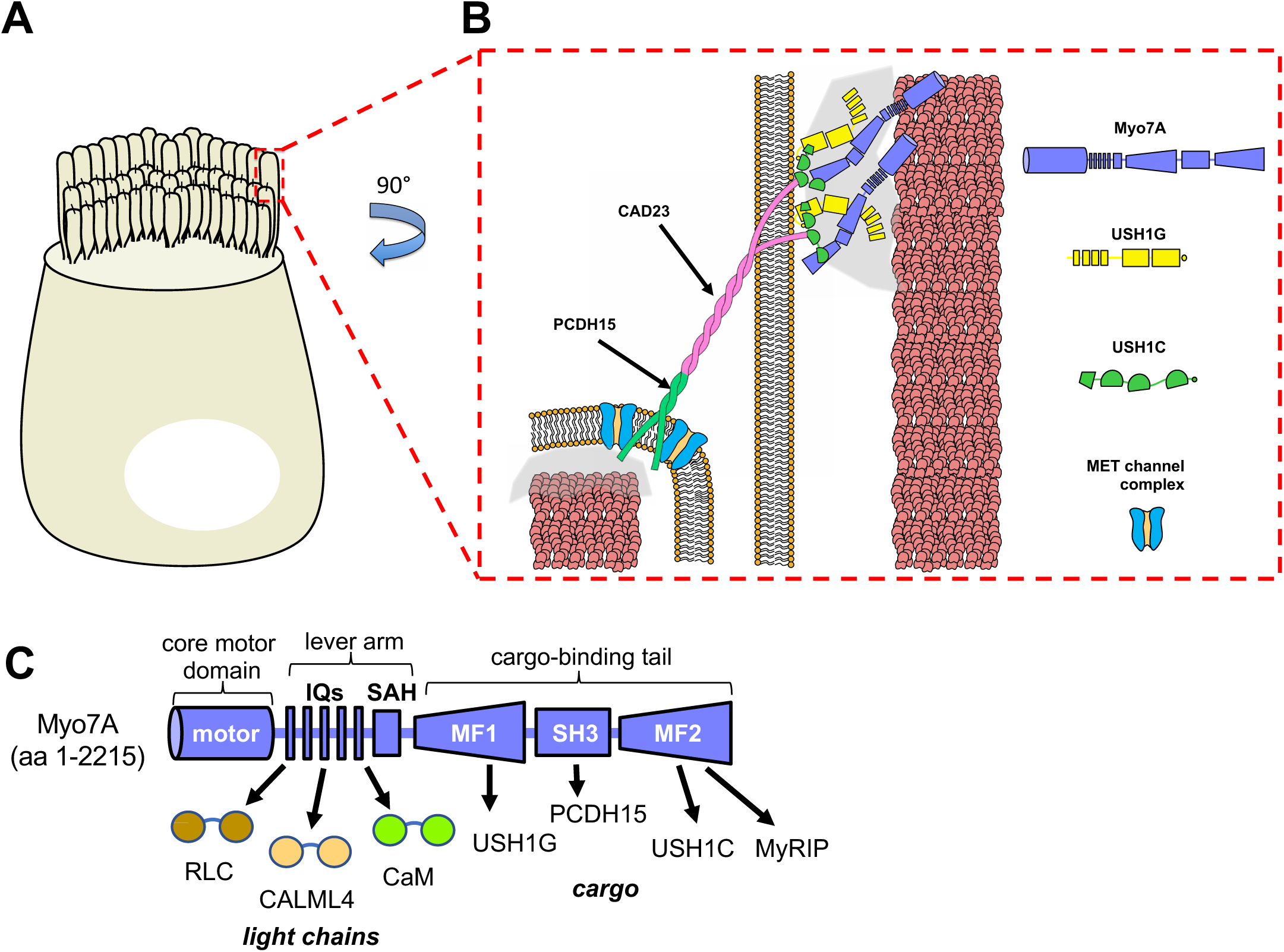
Myo7A in inner ear hair cells. (A) Cartoon diagram of an inner ear hair cell showing a mature hair bundle. (B) Zoom panel showing the tip-link complex that mediates mechanotransduction of sound reception. PCDH15 and CAD23 form the tip-link, with PCDH15 connected to the MET complex and CAD23 forming a complex with Myo7A, USH1G and USH1C. (C) Cartoon domain diagram of Myo7A and its light chains and cargo interaction partners.

Myo7A is composed of an N-terminal core motor domain that binds actin and hydrolyzes ATP in order to convert chemical energy to mechanical force (Fig. 1C). Following the motor domain is a neck region that contains 5 ‘IQ’ motifs that associate with myosin light chains that stabilize the neck, allowing the neck to act as a rigid lever arm during force production. A number of candidate light chains have been identified for Myo7A including conventional calmodulin (CaM), regulatory light chain (RLC), and calmodulin-like protein 4 (CALML4) (Sakai *et al*., 2015; Morgan *et al*., 2016; Choi *et al*., 2020; Hollo *et al*., 2023). The Myo7A lever arm is further extended by a stable single α-helix (SAH) (Yang *et al*., 2009; Peckham, 2011; Li *et al*., 2017). Finally, Myo7A has a C-terminal cargo-binding tail composed of two MyTH4-FERM (MF) domains with an intervening SH3 domain (Fig. 1C). Cargo molecules that associate with the tail include the tip-link components USH1G, USH1C and PCDH15, as well as the Rab-adaptor Myosin and Rab Interacting Protein (MyRIP) (Boeda *et al*., 2002; El-Amraoui *et al*., 2002; Senften *et al*., 2006; Ballesteros *et al*., 2022). Myo7A is thought to exist in an autoinhibited state prior to its activation, in which the cargo-binding tail folds back to contact and inhibit the motor-neck region (Fig. S1) (Umeki *et al*., 2009; Yang *et al*., 2009). Motor inhibition is proposed to be relieved by cargo binding to the tail or by Ca^2+^ binding to the light chains associated with the lever arm (Sakai *et al*., 2011). Cargo is also thought to dimerize Myo7A, creating the motile form of the myosin (Sakai *et al*., 2011; Liu *et al*., 2021). Currently, it is unclear if there are other structural elements of Myo7A that act to dynamically regulate its motor activity in a cellular context. Here, we took the approach of screening for mutations/perturbations to Myo7A that could activate its motor-dependent targeting to the tips of apical microvilli found on the surface of culture kidney proximal tubule epithelial cells. Together, our study provides novel insights into the regulation of Myo7A and how mutations in this important myosin result in sensory disorders in humans.

## RESULTS

### The Myo7A lever arm regulates targeting in CL4 cells

Previous *in vitro* studies of Myo7A have utilized a minimal forced-dimer form of the motor to examine its activity in the absence of cargo binding (Yang *et al*., 2006; Sakai *et al*., 2011; Sakai *et al*., 2015; Sato *et al*., 2017; Matoo *et al*., 2021). This approach involves fusing the isolated motor-neck fragment of Myo7A to a GCN4 leucine zipper dimerization motif in place of its tail (Fig. 2A). The GCN4 motif forces the motor into a dimeric state, promoting a motile form of the myosin without the need for cargo (Fig. 2A). We previously obtained encouraging initial results using this forced-dimer approach in CACO-2_BBE_ enterocytes, where we observed limited targeting of the myosin to the tips of apical microvilli (Matoo *et al*., 2021). However, this cell line has a number of drawbacks that make it a suboptimal model to dissect Myo7A regulation, including a slow doubling time and requiring ∼14-20 days to polarize and assemble mature microvilli. For our current study, we chose to utilize LLC-PK1-CL4 proximal tubule kidney epithelial cells (herein referred to as “CL4 cells”) for a number of reasons. Firstly, Myo7A is natively expressed in proximal tubule cells in kidney tissue (Velichkova and Hasson, 2003). Secondly, CL4 cells rapidly polarize (∼3-4 days) to form well-developed apical microvilli that provide a ‘surrogate’ actin track compared to the actin core found in inner ear stereocilia. Finally, CL4 cells natively express CALML4, an epithelial cell-specific myosin light chain that we previously identified for Myo7A that has recently been confirmed through *in vitro* studies (Choi *et al*., 2020; Hollo *et al*., 2023).

**Figure 2.**
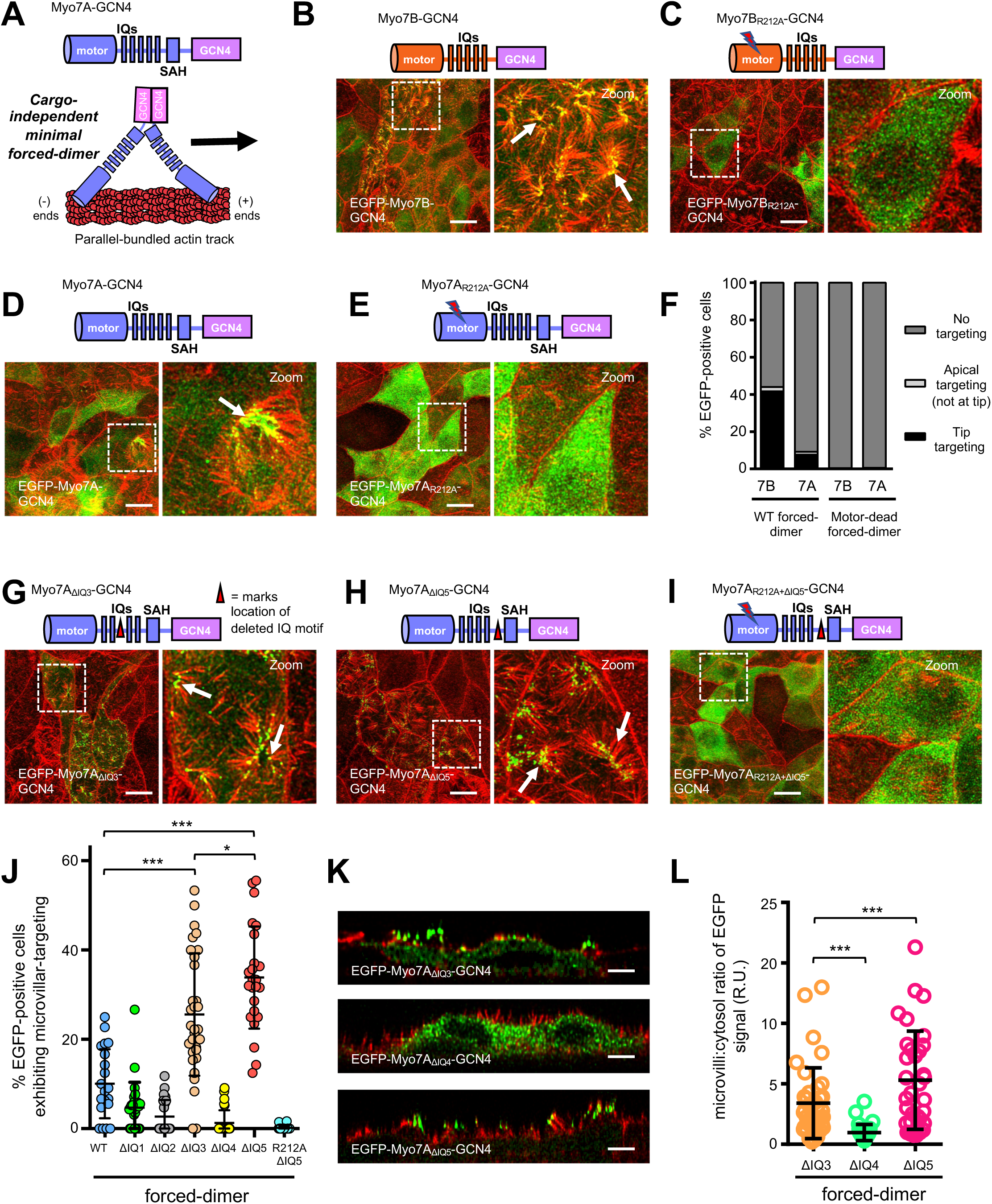
Myo7A targeting activity is controlled by its lever arm in CL4 cells. (A) Cartoon diagram showing the Myo7A forced-dimer. (B-E) Confocal images of CL4 cells stably expressing WT or motor-dead versions of EGFP-tagged Myo7B or Myo7A forced-dimer. Cells were visualized for F-actin (red) and EGFP signal (green). Boxed regions denote areas in zoomed panels for each figure. Arrows point to examples of tip-targeting. Scale bars, 10 µm. (F) Quantification of targeting was performed by binning cells into three categories (no apical targeting, apical targeting but not exclusively at tip, and tip targeting). Tip targeting was then assessed for these cells using line-scan analysis of EGFP signal intensity across microvilli using X-Z sections. Myo7B WT forced-dimer, n = 62 cells; Myo7A WT forced-dimer, n = 65 cells; Myo7B motor-dead forced-dimer, n = 78 cells; Myo7A motor-dead forced-dimer, n = 73 cells. Stacked bars indicate mean. (G-I) Confocal images of CL4 cells stably expressing EGFP-tagged Myo7A forced-dimer containing an internal deletion of IQ3 or IQ5, as well as a motor-dead Myo7A ΔIQ5 forced-dimer control. Cells were visualized for F-actin (red) and EGFP signal (green). Boxed regions denote areas in zoomed panels for each figure. Scale bars, 10 µm. (J) Scatterplot quantification of the percent EGFP-positive cells exhibiting microvillar targeting of the forced-dimer constructs tested. Bars indicate mean and SD. *p < 0.01, ***p < 0.0001, t-test. Bars indicate mean ± SD. (K) Representative confocal cross-section of CL4 stable cell lines expressing EGFP-tagged Myo7A forced-dimer containing an internal deletion of IQ3, IQ4 or IQ5. Cells were visualized for F-actin (red) and EGFP signal (green). Scale bars, 5 µm. (L) Scatterplot quantification of the microvilli:cytosol ratios of EGFP signal for the EGFP-fusion constructs tested. ***p < 0.0001, t-test. Bars indicate mean and SD.

We compared our Myo7A forced-dimer along with an equivalent forced-dimer of Myo7B, the closest homolog to Myo7A, which has been shown to use its motor activity to target to the tips of apical microvilli of CL4 cells (Weck *et al*., 2016). These forced-dimer constructs were stably expressed in CL4 cells as EGFP-fusion proteins and microvillar tip-targeting of the motor quantified by visual inspection of *en face* images of the monolayer. Unexpectedly, while the Myo7B forced-dimer displayed significant tip-targeting in CL4 cells (Fig. 2B,F), the Myo7A forced-dimer exhibited much lower levels of tip-targeting (Fig. 2D,F). We confirmed that microvillar targeting of both forced-dimer constructs were dependent upon their motor activity by incorporating a “motor-dead” mutation (R212A; blocks ATP hydrolysis) into their core motor domains (Fig. 2C,E,F)(Weck *et al*., 2016). These results highlighted that there is a fundamental difference between either the core motor domain or the lever arm (or both) of Myo7A and Myo7B that influences cellular targeting activity between them. Sequence alignments show that the core motor domains of Myo7A and Myo7B exhibit relatively high identity (64%), while the IQ motifs have a much lower identity at 46% (Fig. S2A). To explore whether the Myo7A lever arm plays a role in negatively regulating its targeting activity, we constructed and screened a series of Myo7A forced-dimer constructs in which individual IQ motif were deleted (Fig. S2B). Surprisingly, selective deletion of either IQ3 or IQ5 could significantly activate Myo7A targeting, while deletion of IQ1, IQ2, and IQ4 impaired targeting (Fig. 2G,H,J; Fig. S2C-E). Incorporating the R212A motor-dead mutation into the ΔIQ3 and ΔIQ5 constructs completely blocked targeting, confirming that the targeting was dependent upon a functional motor domain (Fig. 2I,J; ΔIQ5 data shown). Interestingly, we noted that deletion of IQ5 appeared to be a significantly better activator of Myo7A compared to IQ3, with most of the EGFP signal enriched in the microvilli of cells (Fig. 2K). This could be seen by directly comparing the ratio of EGFP signal in the apical microvilli versus the cytoplasm of the Myo7A ΔIQ3 and ΔIQ5 constructs, using ΔIQ4 as a non-activated reference. The ΔIQ5 construct exhibited approximately a ∼2-fold higher enrichment in the apical domain compared to ΔIQ3 (Fig. 2L), suggesting that IQ5 likely functions as the dominant inhibitory IQ motif within the Myo7A lever arm. In contrast, deleting IQ5 from the Myo7B forced-dimer resulted in aberrant localization of the myosin to the nucleus, confirming that the two Class 7 myosins do indeed behave quite differently in cells (Fig. S2F). Together, these results demonstrate that the motor-dependent targeting of Myo7A can be influenced by its lever arm, and that this property does not appear to be strictly shared with its closest homolog Myo7B.

### Point mutations in IQ5 potently activate tip-targeting of Myo7A, including the R853C/H DFNA11 deafness mutants

To further characterize the regulatory role of the Myo7A lever arm, we sought to identify individual point mutations within IQ5 that could replicate the activating effect seen upon deleting the IQ motif. This would allow us to characterize these activating mutants biochemically and examine how they influenced the interaction of IQ5 with its native light chain, CaM. We constructed and screened a series of directed site-saturation mutant libraries covering every amino acid position of IQ5 (Fig. 3A; 20 libraries in total), in order to search for positions that could activate targeting of our Myo7A forced-dimer when mutated. In this brute-force screen, each library represents a pool of EGFP-Myo7A-GCN4 constructs in which a single amino acid position within IQ5 is randomly mutated to any other amino acid (Fig. 3B; see top panel for sequencing example). This screen discovered that mutation of distinct positions within IQ5 promoted robust microvillar targeting of the Myo7A forced-dimer (Fig. 3B). Within IQ5, mutation of positions I851, A852, and R853 (Fig. 3A; residues in cyan) were the most potent activators (Fig. 3B; red line, cutoff above 40% microvillar targeting). Interestingly, these activating positions were clearly biased towards the C-terminal section of IQ5 (Fig. 3B), while mutations in the N-terminal section of IQ5 had little-to-no positive effect on Myo7A targeting activity. We used these libraries to rapidly extract and test defined mutations at each of the most potent amino acid positions identified. We recovered and tested the isolated I851E and A852D mutants (chosen to convert hydrophobic residues to charged side chains), as well as R853C and R853H mutants (chosen since these are reported autosomal dominant DFNA11 deafness mutants)(Bolz *et al*., 2004; Shearer *et al*., 2013; Yamamoto *et al*., 2020). In each case, these mutations promoted robust microvillar targeting of the Myo7A forced-dimer (Fig. 3C-F). Strikingly, the R853C DFNA11 deafness mutation resulted in ∼90% of EGFP-positive cells exhibiting targeting. We further tested other substitutions at R853 and found that loss of a positively charged residue at this position induced the most potent activation of the myosin (Fig. S3A-E).

**Figure 3.**
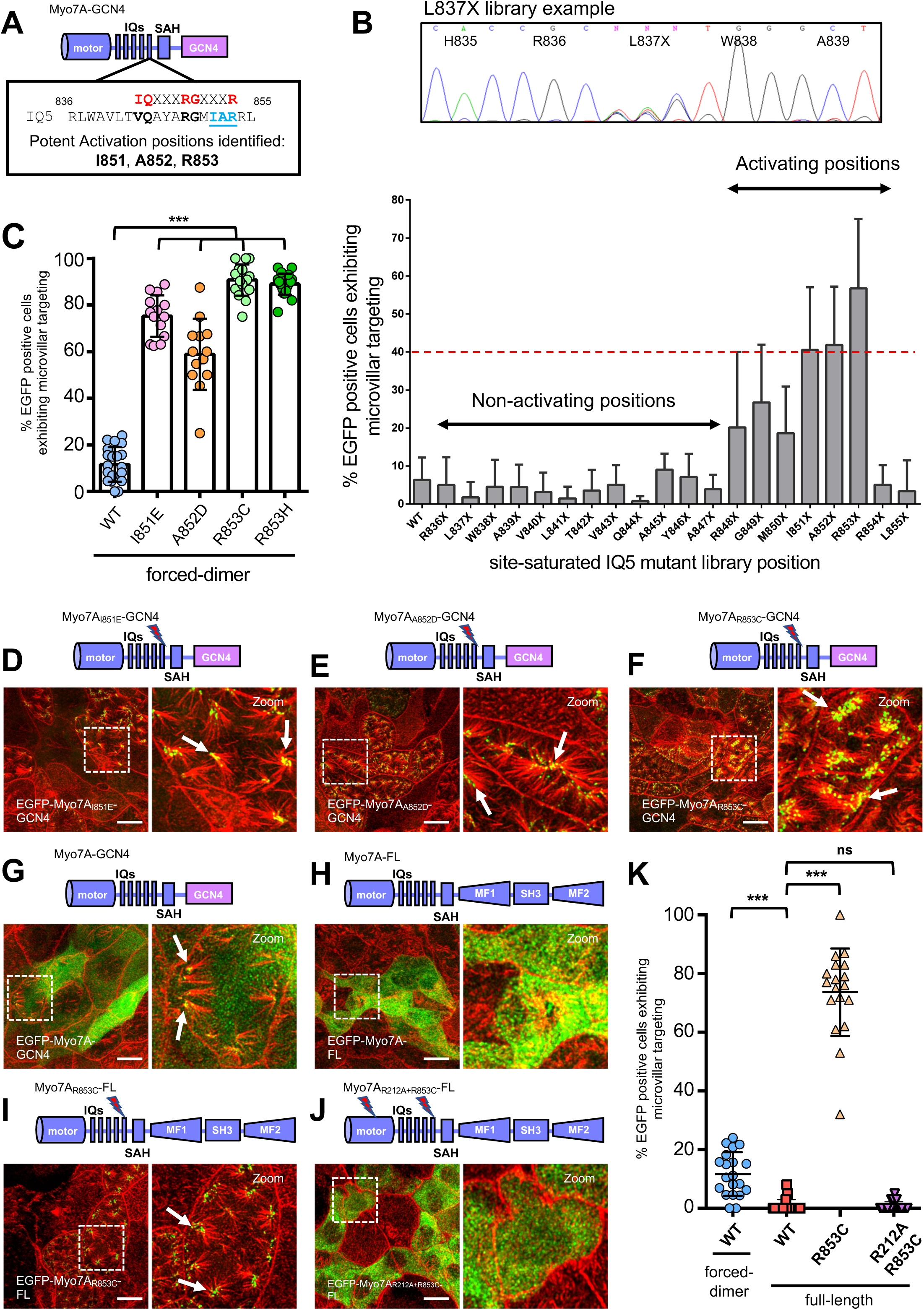
Specific point mutations in IQ5 can activate Myo7A targeting in epithelial cells. (A) Cartoon diagram highlighting the potent activation positions identified in IQ5 of the lever arm of Myo7A. (B) Quantification of microvillar targeting from 20 site-saturated IQ5 mutant Myo7A forced-dimer libraries that were screened in CL4 cells. Red dashed line shows the cut-off of 40% microvillar targeting efficiency that was used to select the most potent activating positions for further analysis. Top panel shows an example of the sequencing of the L837X library. (C) Scatterplot quantification of the percent EGFP-positive cells exhibiting microvillar targeting of the mutant forced-dimer constructs tested. Bars indicate mean and SD. ***p < 0.0001, t-test. Bars indicate mean ± SD. (D-F) Confocal images of CL4 cells stably expressing EGFP-tagged Myo7A mutant forced-dimer. Cells were visualized for F-actin (red) and EGFP signal (green). Scale bars, 10 µm. (G-H) Confocal images of CL4 cells stably expressing EGFP-tagged WT forced-dimer and WT full-length Myo7A. Cells were visualized for F-actin (red) and EGFP signal (green). Scale bars, 10 µm. (I-J) Confocal images of CL4 cells stably expressing EGFP-tagged Myo7A R853C full-length and the corresponding motor-dead control mutant. Cells were visualized for F-actin (red) and EGFP signal (green). Scale bars, 10 µm. (K) Scatterplot quantification of the percent EGFP-positive cells exhibiting microvillar targeting of the WT forced-dimer construct compared to WT full-length and R853C activated full-length. A motor-dead version of Myo7A R853C full-length is also included as a control. Bars indicate mean and SD. ns= not significant, ***p < 0.0001, t-test. Bars indicate mean ± SD.

It is currently proposed that autoinhibition of Myo7A is mediated by its tail domain folding back and contacting the motor domain-lever arm to silence motor activity, and that cargo binding to the tail relieves autoinhibition. Cargo is further thought to oligomerize the motor to promote a motile form of the myosin (Fig. S1). However, our results now suggest that the lever arm alone may be able to act as a direct inhibitor of motor activity, given that our Myo7A forced-dimer construct lacks the tail and appears to be at least partially autoinhibited in cells. To begin to explore whether the lever arm plays an important role in repressing the activity of full-length Myo7A, we questioned whether our IQ5 activating mutations would also promote motor-dependent targeting of full-length Myo7A in CL4 cells. As a starting point, we first compared the levels of microvillar targeting seen between the wild-type (WT) Myo7A forced-dimer and full-length myosin constructs. In contrast to the forced-dimer construct which displayed a low-level of basal targeting (∼15-20%), WT full-length Myo7A exhibited essentially no microvillar targeting in CL4 cells (Fig. 3G,H,K). Incorporation of any of our identified IQ5-activating mutations resulted in robust targeting of full-length Myo7A to the tips of microvilli (Fig. 3I,K; Fig S3F-H,K). This targeting could be ablated by further incorporating the R212A motor-dead mutation (Fig. 3J,K), demonstrating that microvillar tip-targeting was indeed dependent upon the motor activity of the IQ5-activated myosin. Previously, a double point mutation in the second MF domain of Myo7A (R2176A/K2179A; RKAA mutant) was proposed to lock the myosin into an active state by preventing the tail from inhibiting the motor domain (Yang *et al*., 2009; Sakai *et al*., 2015). However, we observed that the RKAA tail mutant was a relatively poor activator of full-length Myo7A in CL4 cells in contrast to our IQ5-actvating mutations (Fig. S3I,K). To assess whether an intact tail domain was necessary for the microvillar targeting ability of our IQ5-activated full-length Myo7A, we truncated part of the C-terminal MF2 domain to disrupt its cargo binding ability and discovered that this significantly reduced tip-targeting (Fig. S3J,K). In sum, these results position IQ5 as being critical in maintaining the inhibited state of Myo7A in cells, and further demonstrate that apical targeting of the IQ5-activated Myo7A in CL4 cells requires both an enzymatically active motor domain and an intact C-terminal cargo-binding tail.

### Activating point mutations in IQ5 block Apo-CaM binding

To investigate how our IQ5-activating mutations influenced light chain binding to the IQ motif, we developed a simple bicistronic E. coli co-expression system to co-express isolated IQ5 fused to GST with CaM light chain in bacterial cells (Fig. 4A). We first confirmed that CaM did not have any non-specific interaction with GST alone in this assay (Fig. S4). While CaM was able to robustly co-purify with GST-7A WT IQ5 both in the presence and absence of Ca^2+^(Fig. 4B; top, left panel), in each case our IQ5-activating mutations specifically blocked calcium-free CaM (herein referred to as Apo-CaM) from binding to IQ5 (Fig. 4B; Fig. S4). This is consistent with previous *in vitro* biochemical studies of the DFNA11 R853C mutation (Bolz *et al*., 2004; Li *et al*., 2017). In combination with our CL4 targeting data of these mutants, this suggests that Apo-CaM bound to IQ5 acts as an inhibitory light chain to silence motor-based targeting of Myo7A in cells.

**Figure 4.**
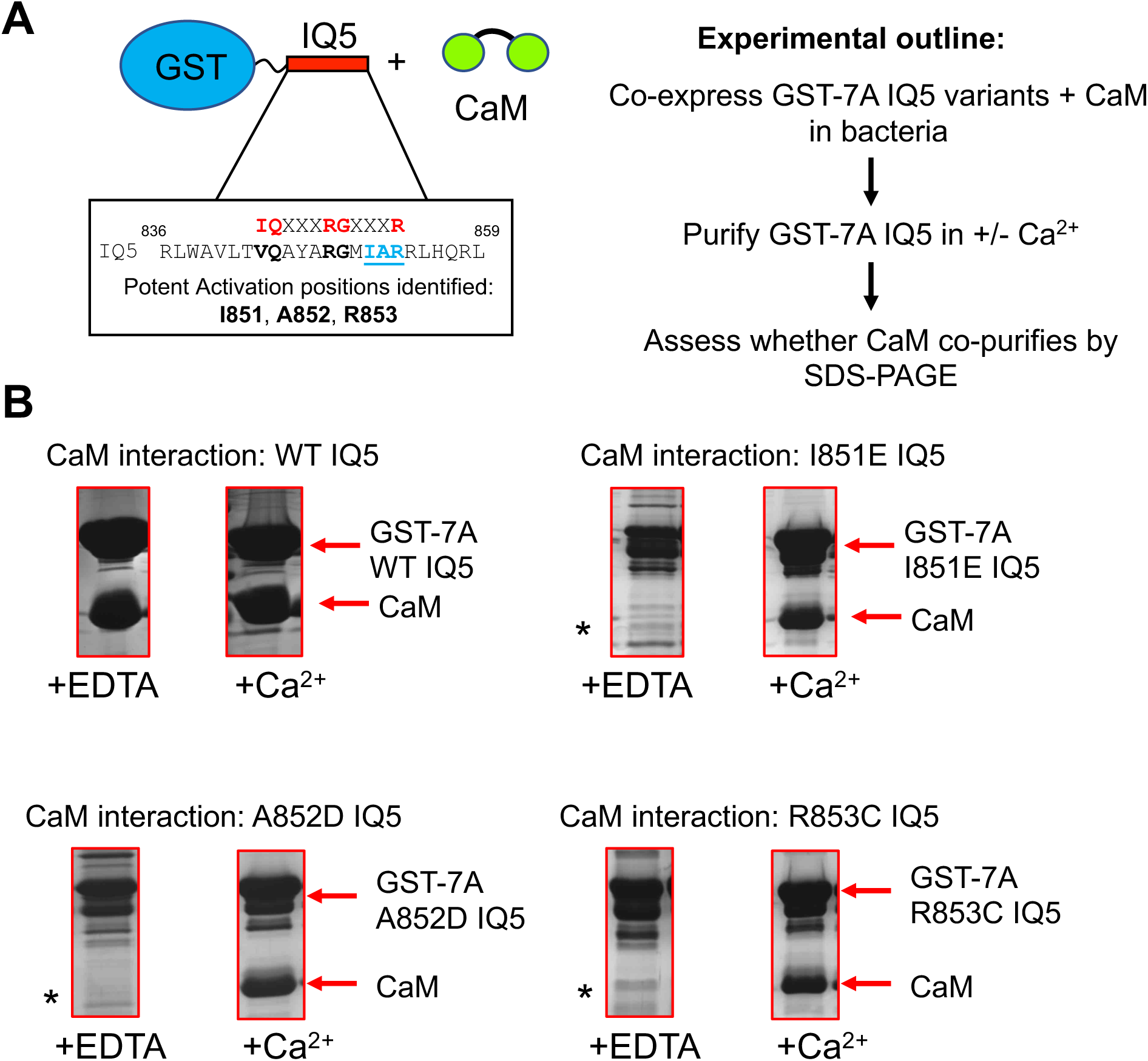
Myo7A-activating mutations found in IQ5 block Apo-CaM from binding to the IQ motif. (A) Cartoon diagram outlining the bicistronic E. coli expression system used to test the interaction between IQ5-activating mutants and CaM. (B) Coomassie-blue stained SDS-PAGE gels showing the pull-down results between IQ5-activating mutants and CaM done in the presence and absence of Ca^2+^. WT IQ5 is shown as a positive control. Migration of the GST-IQ5 and CaM protein are shown by arrows, while the stars (*) denote the loss of binding of the IQ5 mutant with Apo-CaM.

### Calcium does not regulate Myo7A targeting in cells

Early studies of recombinant Myo7A have suggested that Ca^2+^ binding to the light chains associated with the lever arm could relieve tail-mediated autoinhibition (Umeki *et al*., 2009; Yang *et al*., 2009). Our discovery that mutations that block Apo-CaM from binding to IQ5 potently activate Myo7A targeting immediately raised the possibility that Ca^2+^ could regulate Myo7A activity, by converting Apo-CaM to Ca^2+^-CaM. To investigate this possibility, we utilized the Ca^2+^ ionophore drug A23187 to artificially raise cytosolic levels of Ca^2+^ to determine whether this could act as a temporal signal to activate Myo7A in cells. As proof-of-principle, we first tested whether Ca^2+^ ionophore A23187 could, indeed, increase cytoplasmic levels of Ca^2+^ in CL4 cells. We generated a stable CL4 cell line expressing the widely-used GCaMP6 genetically-encoded Ca^2+^ sensor protein that fluoresces when bound to calcium (Chen *et al*., 2013) to use as a readout of cytosolic calcium levels. Live-cell imaging of this stable cell line revealed a rapid increase in fluorescence signal from cells upon exposure of Ca^2+^ ionophore A23187 (Fig. 5A; Video S1). However, despite trying numerous concentration ranges and time courses, we found that treatment of our CL4 EGFP-Myo7A full-length stable cell line with Ca^2+^ ionophore resulted in only a minor (but not statistically significant) increase in Myo7A targeting (Fig. 5B-F; Fig S5A,B). This data demonstrates that cytosolic Ca^2+^ levels likely do not function as a major regulator of Myo7A in cells, suggesting other mechanisms likely governs its activity.

**Figure 5.**
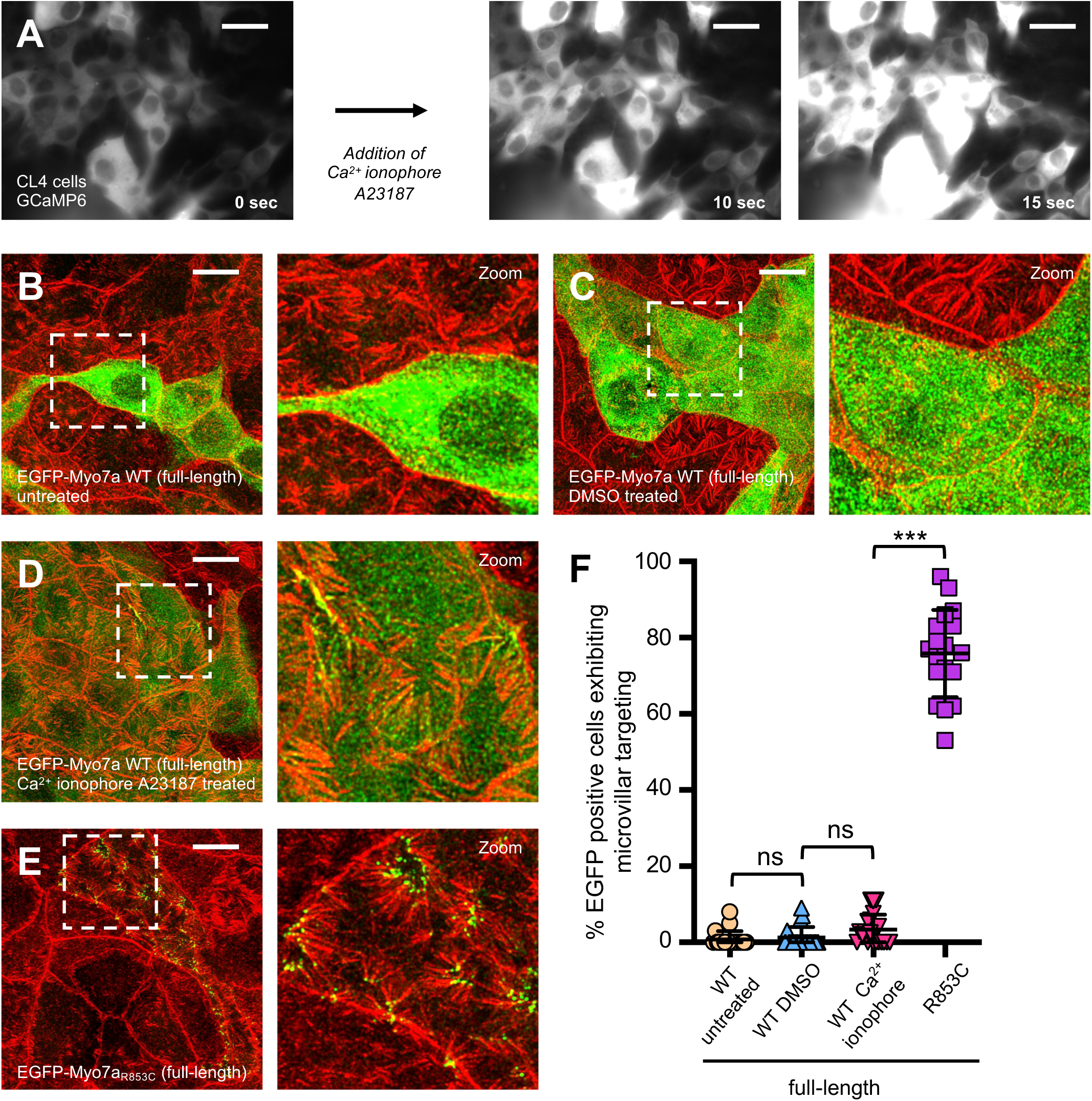
Raising intracellular Ca^2+^ levels does not result in potent activation of Myo7A targeting in CL4 cells. (A) Live-cell images of CL4 cells stably expressing the genetically-encoded Ca^2+^ sensor protein GCaMP6 that fluoresces when bound to calcium. Exposure to 1µm Ca^2+^ ionophore A23187 resulted in a rapid increase in GCaMP6 fluorescence, indicating increased cytoplasmic Ca^2+^ levels. Scale bars, 25 µm. (B-E) Confocal images of CL4 cells stably expressing EGFP-tagged Myo7A WT full-length construct that were exposed to either DMSO alone or 1µm Ca^2+^ ionophore A23187. EGFP-tagged Myo7A R853C full-length construct was used as a control to show the level of microvillar targeting achieved with genetic activation of the myosin. Cells were visualized for F-actin (red) and EGFP signal (green). Scale bars, 10 µm. (F) Scatterplot quantification of the percent EGFP-positive cells exhibiting microvillar targeting of EGFP-tagged Myo7A full-length cell lines treated with DMSO alone or 1µm Ca^2+^ ionophore A23187. Microvillar targeting of the EGFP-tagged Myo7A R853C full-length construct was used as a comparison. Bars indicate mean and SD. ns=not significant, ***p < 0.0001, t-test. Bars indicate mean ± SD.

### Select DFNA11 mutations found in the core motor domain and SAH motif of Myo7A activate targeting

We expanded our analysis to determine whether other reported DFNA11 mutations that lie outside the Myo7A IQ motifs had a similar effect on the targeting of the myosin in CL4 cells. We first tested 15 reported DFNA11 variants found in the core motor domain by incorporating these mutations into the context of the Myo7A forced-dimer (Fig. 6A) (Luijendijk *et al*., 2004; Street *et al*., 2004; Bischoff *et al*., 2006; Di Leva *et al*., 2006; Kallman *et al*., 2008; Su *et al*., 2009; Sun *et al*., 2011; Sang *et al*., 2013; Iwasa *et al*., 2016; Kaneko *et al*., 2017; Li *et al*., 2018; Lu *et al*., 2020; Joo *et al*., 2022; Kim *et al*., 2023; Xia *et al*., 2023) (Supplemental Table 1). For this analysis, we normalized the targeting activity of all constructs against WT forced-dimer levels. Of these 15 core motor domain mutations, 8 were found to significantly activate Myo7A targeting (Fig 6B). Mapping these activating positions onto an available structural model of Myo7A (Kuppa and Sergeev, 2021) revealed that they cluster into two general areas of the core motor; a flat broad surface of the motor equivalent to the region originally described as the myosin mesa in studies of β-cardiac myosin, and secondly the interface between where the relay and SH1 helices abut against the converter domain (Fig. 6C; Fig. S6A,B). Within the myosin mesa, the most potent activator was the D218N mutation which impacts a solvent-facing aspartic acid residue found as part of one of the β-strands of the transducer β-sheet (Fig. 6D). Activating mutations that mapped to the relay helix-converter domain interface were found on both sides of this interface; for example, R668H impacts the SH1 helix that leads into the converter domain (Fig. 6E), while R675C changes a highly conserved arginine residue in the small β-sheet of the converter domain that abuts against the relay helix (Fig. 6F). In each case, we found that these relay helix-converter domain DFNA11 mutations were able to potently activate targeting of Myo7A in our CL4 cells. We further tested the DFNA11 motor domain mutations in the context of full-length Myo7A, as well as the mutations reported in the SAH motif and tail domain. Nearly all the DFNA11 mutations that activated the forced-dimer construct were able to activate full-length myosin (Fig. S6C,D; targeting of full-length R675C shown as example). The notable exceptions were R616Q and R623H found in the myosin mesa that were able to partially activate the forced-dimer, but had no effect on full-length targeting, and the relay helix mutation L479P and G660R mutation found in the connection between the SH2 and SH1 helices, that did not significantly activate the forced-dimer, but were able to activate full-length Myo7A. We also observed that a DFNA11 variant that results in a three amino acid deletion in the SAH motif (Δ886-888) was able to activate full-length Myo7A targeting to levels near our IQ5-activating mutants (Fig. S6C,E). In stark contrast, none of the DFNA11 mutations reported in the tail domain of Myo7A activated its targeting in CL4 cells.

**Figure 6.**
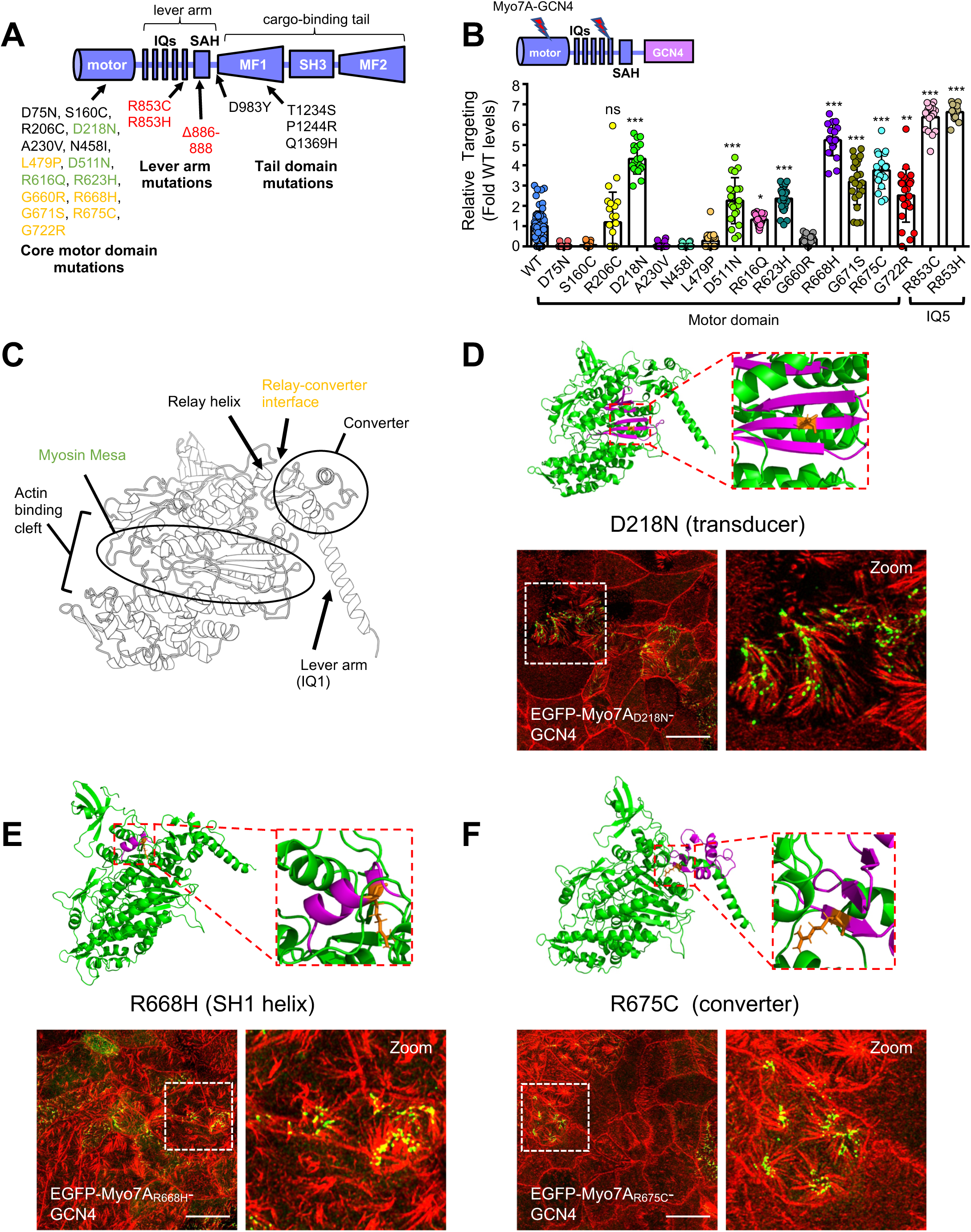
Select DFNA11 mutations found in the core motor domain and SAH motif activate Myo7A targeting. (A) Cartoon diagram showing the location of DFNA11 mutations that have been investigated in our study. Variants in green map to the myosin mesa, variants in yellow map to the relay helix-converter domain interface, and variants in red map to the lever arm. (B) Scatterplot quantification of the relative microvillar targeting of the DFNA11 variants compared to WT Myo7A forced-dimer. All mutants were tested in the context of the Myo7A forced-dimer construct. Bars indicate mean and SD. ns= not significant, *p < 0.01, **p < 0.001, ***p < 0.0001, t-test. Bars indicate mean ± SD. (C) Model of Myo7A highlighting what structural features are impacted by DFNA11 motor domain variants that activate targeting of the myosin in CL4 cells. (D-F). Examples of DFNA11 motor domain variants that activate targeting of the myosin in CL4 cells. Shown are confocal images of CL4 cells stably expressing each EGFP-tagged Myo7A forced-dimer mutants. Cells were visualized for F-actin (red) and EGFP signal (green). Scale bars, 10 µm. Above each confocal image set are structural models showing what structural element (magenta) is impacted by the particular DFNA11 mutation (residue mutated is shown in orange stick form).

We noted that a number of the non-activating DFNA11 motor domain mutations actually appeared to abolish the basal level of apical targeting of our Myo7A forced-dimer construct (Fig. 6B; D75N, S160C, A230V, N458I), suggesting that they may actually inhibit rather than promote motor activity of Myo7A. To explore this further, we incorporated these mutations into the context of our Myo7A forced-dimer that was activated using the R853C IQ5 mutation, and quantified the level of targeting of the double mutants in comparison to the R853C mutant alone (Fig. S6F). As a control, we also tested similar double mutants using two motor domain variants that we discovered potently activated Myo7A targeting (D218N and R675C) (Fig. S6F). While D218N and R675C had neither a positive or negative effect on the targeting of IQ5-activated Myo7A, the other select mutations tested did indeed impair the targeting of the IQ5-activated Myo7A forced-dimer, suggesting that they likely compromise the motor output of the myosin. Together, our data demonstrate that the majority of reported DFNA11 deafness variants reported in the motor domain and lever arm of Myo7A have the ability to activate targeting of the myosin in cultured epithelial cells.

### IQ5-activated Myo7A mistargets in inner ear hair cells

Our discovery that select DFNA11 mutations constitutively activate targeting of Myo7A in our kidney cell culture model suggests that the molecular basis of DFNA11 deafness may be due to dysregulation of the myosin, rather than having a motor with impaired enzymatic function. To begin to shed light on the etiology of DFNA11 deafness, we examined targeting of our IQ5-activated Myo7A directly in inner ear hair cells. Inner ear hair cell tissue explants were harvested from C57BL/6 mice (P5 stage) and transfected with either EGFP-tagged WT or R853C mutant full-length constructs of Myo7A. WT Myo7A typically exhibited little targeting, with only rare occurrences in which it was found enriched in stereocilia (Fig. 7A; Fig. S7A; targeting examples shown). For WT transfected cells that did appear to have signal localized in stereocilia, line-scan analysis across the rows of stereocilia revealed that most of the EGFP signal was enriched in the short and middle stereocilia rows, where endogenous Myo7A is known to be found (Fig. 7B) (Grati and Kachar, 2011). In contrast, IQ5-activated Myo7A R853C was mislocalized to the distal tips of the longest row of stereocilia in almost all successfully transfected cells (Fig. 7C,D; Fig. S7B). Together with our other studies, these data suggest that autosomal dominant DFNA11 deafness mutations dysregulate Myo7A, likely causing it to aberrantly localize in inner ear hair cells because the myosin is constitutively active.

**Figure 7.**
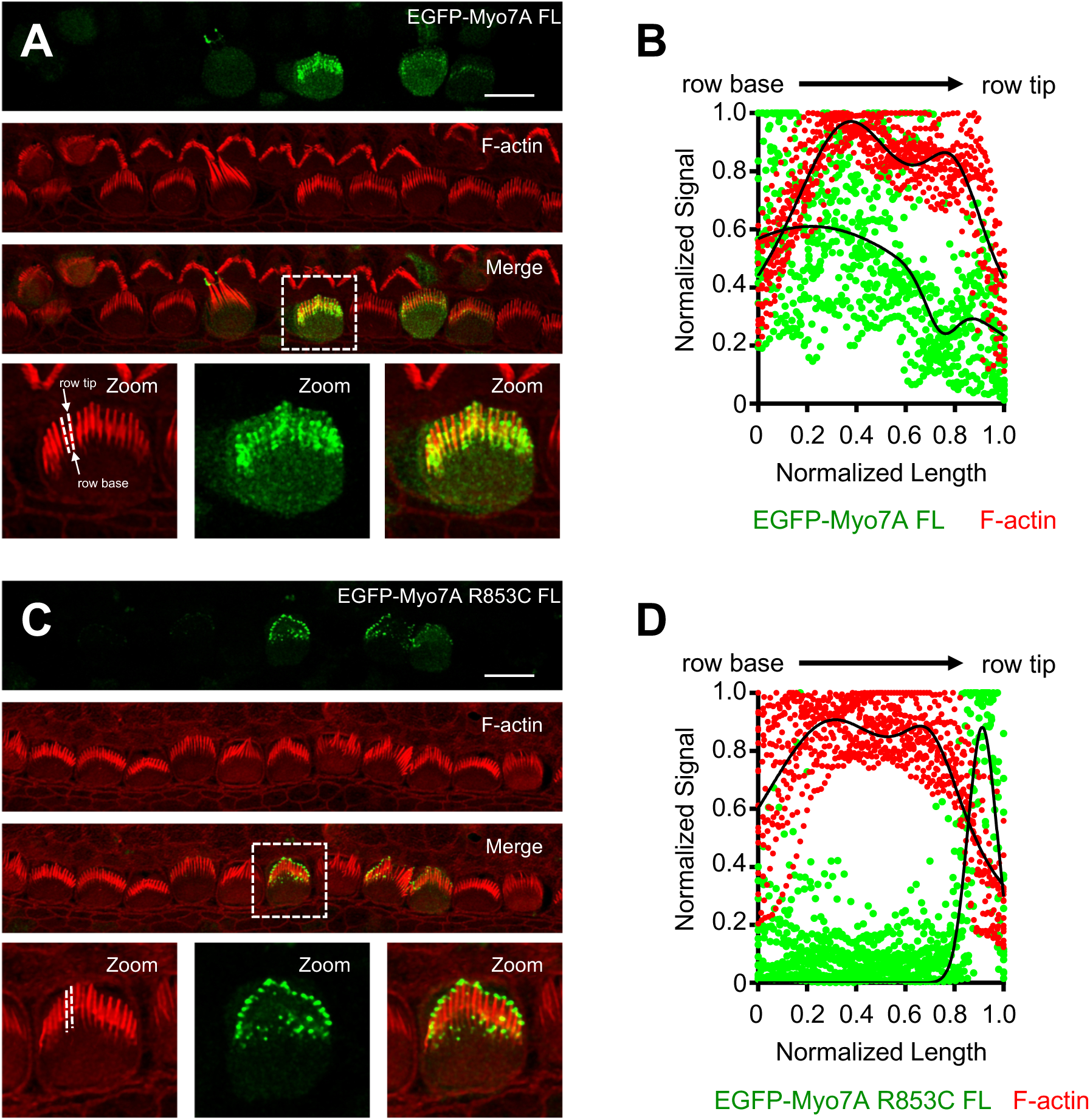
The R853C DFNA11 mutation causes Myo7A to aberrantly target in mouse explant cultured inner ear hair cells. (A) Confocal images of mouse tissue explant cultured inner ear hair cells transfected with EGFP-tagged Myo7A WT full-length. Shown is an example in which the WT Myo7A exhibited targeting to stereocilia. Cells were visualized for F-actin (red) and EGFP signal (green). White dashed lines in the F-actin zoom panel provide examples of how line-scans were performed. Scale bar, 10 µm. (B) Line-scan analysis across the rows of stereocilia for WT transfections that appeared to localize in stereocilia. EGFP signal has been normalized across all scans, as well as the lengths of the line-scan where 0=row base and 1=row tip. (C) Confocal images of mouse tissue explant cultured inner ear hair cells transfected with EGFP-tagged Myo7A R853C full-length. In contrast to WT, almost all successful R853C transfections had signal for the myosin in stereocilia. Cells were visualized for F-actin (red) and EGFP signal (green). White dashed lines in the F-actin zoom panel provide examples of how line-scans were performed. Scale bar, 10 µm. (D) Line-scan analysis across the rows of stereocilia for R853C transfections. EGFP signal has been normalized across all scans, as well as the lengths of the line-scan where 0=row base and 1=row tip.

## DISCUSSION

The first autosomal dominant DFNA11 variant discovered was mapped to cause a three amino acid deletion (Δ886-888) in the Myo7A protein, impacting what was originally thought to be a coiled-coil motif in the lever arm/proximal tail region (Tamagawa *et al*., 1996; Weil *et al*., 1996; Liu *et al*., 1997). At the time, this predicted coiled-coil motif was proposed to function as an inherent dimerization motif for the myosin, since a pairwise yeast-two-hybrid experiment utilizing a fragment encoding the predicted coiled-coil was positive for homomeric interaction (Weil *et al*., 1997). However, subsequent chemical cross-linking and electron microscopy studies of purified recombinant human and Drosophila Myo7A did not detect evidence of inherent dimerization, and the predicted coiled-coil was later shown to function as a SAH motif instead (Umeki *et al*., 2009; Yang *et al*., 2009; Li *et al*., 2017). More recent mass photometry experiments agree that Myo7A, on its own, appears to be a monomer (Liu *et al*., 2021; Hollo *et al*., 2023). The original idea that Myo7A contained an inherent dimerization motif, coupled with the discovery of an autosomal dominant deafness mutation in this motif, lead to the proposal that DFNA11 Myo7A variants might be pathogenic by acting as enzymatically-dead dominant negative copies of the myosin that dimerize with WT protein in cells (Liu *et al*., 1997; Bolz *et al*., 2004). In this manner, the molecular basis of DFNA11 deafness was thought to be similar in nature to DFNB2 and USH1B patients in that DFNA11 patients would also have a “deficit” of Myo7A activity. Our results now suggest a much more likely scenario in which DFNA11 mutations constitutively activate the myosin, providing a simple explanation of the autosomal dominant genetics of DFNA11 deafness.

Early biochemical and biophysical studies utilizing purified recombinant Myo7A lead to the model in which the tail folds back upon the motor domain to regulate the activity of the myosin (Umeki *et al*., 2009; Yang *et al*., 2009). Cargo binding to the tail or Ca^2+^ binding to light chains associated with the lever arm were mechanisms proposed to relieve this intramolecular autoinhibition. Importantly, these early studies also identified a critical role for the distal IQ motifs in regulating Myo7A; an isolated fragment encoding the tail of Myo7A could inhibit the ATPase activity of the motor domain in *trans* only if the motor domain fragment also included the distal IQ motifs (Umeki *et al*., 2009). Our studies here can now further refine this model. We identify IQ3 and IQ5 as the regulatory IQ motifs within the Myo7A lever arm and show that Apo-CaM bound to IQ5 is inhibitory. Despite the apparent importance of the IQ5/CaM interaction in controlling Myo7A activity in cells, elevating intracellular Ca^2+^ levels did not appear to be able to activate the myosin in cells. In agreement with this, recent *in vitro* motility assays performed by the Liu and Sellers labs demonstrated that elevated Ca^2+^ concentrations actually hamper the ability of purified Myo7A to move actin filaments, indicating a compromised mechanical function of the motor (Hollo *et al*., 2023). Together, these results point against the idea that cellular Ca^2+^ levels can positively regulate Myo7A activity *in vivo*. Perhaps one of the more interesting discoveries from our study is that we can potently activate motor-dependent targeting of full-length Myo7A in CL4 cells using single point mutations in IQ5, but this targeting requires an intact MF2 cargo-binding tail domain. Although we cannot rule out other possible scenarios, this may suggest that CL4 cells express endogenous cargo for Myo7A that can support its motility (possibly through dimer formation), but this cargo on its own cannot activate the myosin. This suggests that there could be different “classes” of cargo for Myo7A: those that activate the myosin, those that support its motility, or those that can do both functions. Although not the subject of this study, we did confirm that both of the tip-link scaffold proteins (USH1C and USH1G) are expressed in CL4 cells (Acharya, unpublished results), which indicates that these cargoes on their own cannot efficiently activate full-length Myo7A in cells. Finally, we also observed that our Myo7A forced-dimer construct, which lacks the tail, appeared to be partially autoinhibited in cells. Previous studies using forced-dimer approaches to study Myo7A in cells have largely assumed that this form of the myosin would be fully active, which might not be the case. Indeed, in our experience caution is warranted when designing artificial means of dimerizing a minimal motor-neck fragment of Myo7A. We observed that the nature of the dimerizing modality can influence the basal level of targeting of a WT Myo7A forced-dimer construct (Acharya et al., manuscript in preparation). For example, we saw using the FKBP inducible homodimerization tag (∼12kDa) resulted in significantly higher levels of basal targeting compared to using the smaller GCN4 motif (3.8 kDa) as the dimerizer. When fused to the C-terminus of a Myo7A motor-neck fragment, a larger dimerizing tag may sterically block the lever arm from inhibiting the motor domain, in contrast to a smaller tag. Future structural studies of full-length Myo7A will be required to resolve how exactly the lever arm and the tail domain contribute to the nature of the autoinhibited state of Myo7A.

Along with the lever arm, we identify specific DFNA11 mutations directly in the motor domain of Myo7A that can activate targeting of the myosin in cells. These motor domain mutations map to two general areas of the motor: the myosin mesa, and the interface between the relay helix and the converter domain. Interestingly, mutations in these two structural areas in β-cardiac myosin are known to cause autosomal dominant hypertrophic cardiomyopathy (HCM) (Robert-Paganin *et al*., 2018; Spudich *et al*., 2024). Similar to our DFNA11 mutations in Myo7A, autosomal dominant HCM mutations are thought to aberrantly activate β-cardiac myosin. The emerging view is that HCM mutations largely prevent β-cardiac myosin from forming an autoinhibited state, part of which involves its motor domain folding back upon its proximal S2 sequence in the inhibited dimer state (Grinzato *et al*., 2023). During our analysis of Myo7A, it did not escape our attention that a number of DFNA11 motor domain mutations that potently activate Myo7A in our cell system (D218N, myosin mesa; R675C, converter-relay helix interface) map to identical structural positions at the residue level that are mutated in HCM (β-cardiac myosin: R249Q/N, myosin mesa; R712L, converter-relay helix interface). It is also interesting to note that the SAH motif of Myo7A is similar in nature to the proximal S2 sequence of β-cardiac myosin. Both of these elements are variations of straight rod-like structures that are made up of alternating regions of negatively and positively charged residues. It was originally proposed that the SAH motif of Myo7A simply functions to extend the working stroke of its lever arm (Baboolal *et al*., 2009; Li *et al*., 2017). We discovered that the Δ886-888 DFNA11 variant found in the SAH motif of Myo7A was able to potently activate tip-targeting of the myosin. This may suggest that the Myo7A SAH motif could play a more complex role in the myosin, possibly contributing important interactions towards forming the autoinhibited state of Myo7A. In agreement with this, a recent cryo-EM structure of Myo6 identified a unique SAH-extension sequence that directly interacts with the active site of its motor domain to promote the autoinhibited state of this myosin (Niu *et al*., 2024). All together, these observations raise the intriguing possibility that even physiologically divergent myosins such β-cardiac myosin and Myo7A may share similar structural themes in how their motor activities are regulated *in vivo*. This may be important given that small molecule modulators of β-cardiac myosin have been developed to treat HCM, suggesting that a similar treatment strategy could be a viable option to pursue for the treatment of DFNA11 deafness.

## ACKNOWLEDGMENTS

We thank all members of the Crawley laboratory for advice and support. This work was supported by a University of Toledo Graduate Student Association Award (P.A.), Northern Ohio Alliances for Graduate Education and the Professoriate (NOA-AGEP) fellowship (MJG), Undergraduate Summer Research and Creative Activities Program (USR-CAP) University of Toledo scholarships (A.T.K., and A.M.), University of Toledo Academic Year Research Awards (FA, TN and JM), NIH R01DC015495 (BJP), and NIH R15GM131382, R15CA264735 awards (SWC).

## AUTHOR CONTRIBUTIONS

PA, GT, XL, SM, MJG, SYA, AKT, TN, NMC, JM, MRH, FA, AR, AM, JB, NR, KH, VN, RA and SWC data curation; PA, GT, and SWC formal analysis; PA, GT, and SWC investigation; PA, BJP, SWC writing-editing; SWC conceptualization; BJP and SWC supervision; BJP and SWC funding acquisition; SWC methodology; SWC writing-original draft; SWC project administration.

## MATERIALS AND METHODS

### Molecular biology

The human cDNA constructs used in this study are as follows: Myo7A, GI: 1519245357, UniProtKB-Q13402; Myo7B, GI: 122937511, UniProtKB-Q6PIF6. DNA encoding these components were generated by PCR and cloned into the pCR8 or the pDONR entry vectors (Invitrogen). Verification of all constructs were performed by sequencing. The resulting entry vectors were then shuttled into destination expression vectors using the LR Clonase II enzyme. Domain boundaries and the design of constructs were aided using information from UniProt. Motor-neck fragments of Myo7A and Myo7B were fused to a GCN4 leucine zipper motif to create forced-dimer constructs. The domain boundaries for Myo7A constructs used are follows: Myo7A WT forced-dimer (aa 1-1014), Myo7A_ΔIQ1_ forced-dimer (aa 1-1014, Δ742-766), Myo7A_ΔIQ2_ forced-dimer (aa 1-1014, Δ768-788), Myo7A_ΔIQ3_ forced-dimer (aa 1-1014, Δ791-811), Myo7A_ΔIQ4_ forced-dimer (aa 1-1014, Δ812-836), Myo7A_ΔIQ5_ forced-dimer (aa 1-1014, Δ837-857), Myo7A full-length (aa 1-2215), Myo7A_Δ886-888_ full-length (aa 1-2215, Δ886-888), Myo7A ΔC-term (aa 1-2175). The domain boundaries for Myo7B constructs used are follows: Myo7B WT forced-dimer (aa 1-968), Myo7B_ΔIQ5_ forced-dimer (aa 1-968, Δ857-884). All point mutations and amino acid deletions were incorporated into constructs using the QuikChange site-directed mutagenesis kit (Agilent). All constructs were expressed using either the pINDUCER20-C1-EGFP lentiviral vector (Crawley *et al*., 2016) or the pEGFP-C1-GW vector (Matoo *et al*., 2024) that have an EGFP-tag upstream of the Gateway recombination cassette. These vectors were used for the generation of stable cell lines expressing EGFP-fusion proteins. Bicistronic E. coli expression constructs were designed by first fusing the isolated Myo7A IQ5 motif (aa 830-859) in-frame to GST in the pGEX-4T3 vector using BamHI-XhoI sites. Subsequently, a synthetic gene fragment encoding CaM along with an upstream ribosome binding site sequence was produced (Twist Biosciences) and cloned downstream into the pGEX-4T3-Myo7A IQ5 vector using XhoI-NotI restriction sites. As a control, the rbs-CaM gene fragment was cloned into the empty pGEX-4T3 vector to test the control interaction between GST alone and CaM. Point mutants of IQ5 were generated using the QuikChange site-directed mutagenesis kit (Agilent).

### Directed site-saturation mutant libraries

Directed site-saturation mutant libraries were created using a modified version of the Myo7A forced-dimer construct cloned into the pEGFP-C1 vector, designed to have a removable fragment of the lever arm that encodes part of IQ3, all of IQ4 and IQ5, and part of the SAH (amino acids 805-868). This fragment could then be replaced with directed site-saturation mutant library fragments that were synthesized. Specifically, this Myo7A forced-dimer template was created by engineering a silent HindIII site at amino acids 805-806 (RSR**KL**HQQ; tail end of IQ3) and the native HindIII site was deleted at amino acids 879-880 (EEE**KL**RKE; middle of SAH). The amino acid 805-865 fragment of the lever arm could then be removed by cutting the engineered HindIII and using a native XhoI site at amino acids 868-869 (YLWR**LE**AEK; beginning of SAH). 20 individual synthetic DNA lever arm fragment libraries were synthesized based on this swappable lever arm fragment and cloned HindIII-XhoI into the pTWIST-AMP vector (Twist Biosciences). Each library represents a lever arm fragment that has a single position within IQ5 mutated to a random amino acid. These synthetic libraries are digested from pTWIST-AMP vector and ligated into pEGFP-C1-Myo7A GCN4 vector and then subsequently transfected into LLC-PK1-CL4 cells using PEI transfection for stable cell line selection.

### Cell culture, lentivirus production, and stable cell line generation

LLC-PK1-CL4 and HEK293FT cells were cultured at 37°C and 5% CO2 in Dulbecco’s modified Eagle’s medium with high glucose and 2mM L-Glutamine (Sigma-Aldrich). 10% fetal bovine serum (FBS) was added to medium for both LLC-PK1-CL4 and HEK293FT cells. Lentivirus particles were generated by co-transfecting HEK293FT cells (10 cm dish at 80% confluency) with 6 μg of lentiviral overexpression plasmid with 4 μg psPAX2 packaging plasmid (Addgene #12260) and 0.8 μg pMD2.G envelope plasmid (Addgene #12259) using polyethylenimine reagent (Polysciences) for Myo7A forced-dimer constructs or Lipofectamine (Lipofectamine 2000) for Myo7A full-length constructs. Cells were incubated with transfection medium for 12 hrs, after which they were exchanged with fresh medium and subsequently incubated for 2 days to allow production of lentivirus into the medium. Medium containing lentiviral particles was collected and filtered with a 0.45 μm syringe filter and concentrated with the addition of Lenti-X concentrator reagent (Takara). For lentiviral transduction, LLC-PK1-CL4 cells were grown to 90% confluency in T25 flasks. 8 μg/ml polybrene (Sigma-Aldrich) was added to the medium after which the cells are transduced with ∼300 μl of concentrated lentiviral particles. For pEGFP-C1-vector transfections, LLC-PK1-CL4 cells were grown to 90% confluency in T25 flasks and incubated with transfection mixture of using 4 μg DNA with polyethylenimine reagent. After 12 hrs of incubation with lentivirus or transfection mixture, the cells ere reseeded into 10 cm dishes and grown for 3 days. Cells were then reseeded into T182 flasks with medium supplemented with 1 mg/ml G418 (Santa Cruz Biotechnology) and passaged to select for stable integration. Induction of pINDUCER20 EGFP constructs was driven by adding 1 μg/ml doxycycline in the medium.

### Protein purification

Bicistronic expression plasmids were transformed into the T7 express E. coli cell line and grown to an O.D =1 at 37°C, after which the temperature was lowered to 24°C and expression induced overnight (12-16 hrs) using IPTG, 1mM (Invitrogen). Cells were then harvested by centrifugation at 4000 Ξ g for 20 mins and purified GSH resin (Sigma-Aldrich) using standard conditions. Briefly, cells were lysed using cold lysis buffer (1XPBS, pH 7.4, 1mg/ml lysozyme, 1mM PMSF, one Protease inhibitor tablets/40ml of Lysis buffer) along with sonication. The cell lysate was then centrifuged for 1 hr at 15000 X g and the supernatant recovered and divided into two equal volumes in chilled conical tubes, in which one received 2mM Ca^2+^ and the other 2mM EDTA. These lysates were then incubated with GSH resin on a rocking platform for 1 hr at 4°C. The lysate-GSH resin mixture was added to a column, washed with Wash Buffer (1XPBS pH 7.4) and the GST fusion protein eluted with Elution Buffer (1XPBS pH 7.4, 20mM Glutathione). Eluates were analyzed by SDS-PAGE.

### Microscopy

LLC-PK1-CL4 stable cell lines were seeded on coverslips at 80% confluency and allowed to grow/polarize for 4 days. Cells were then fixed in 4% paraformaldehyde (Electron Microscopy Sciences) in PBS for 15 mins at RT, washed with PBS, and permeabilized with 0.1% Triton X-100 (Sigma) in PBS for 7 mins. After fixation, cells were washed four times with PBS and then blocked with 5% BSA overnight at 4°C. Staining was performed with Alexa Fluor 568 phalloidin (1:200) at RT for 1 hr. The coverslips were then washed four times in PBS and then mounted using ProLong Diamond Anti-fade reagent (Invitrogen). Cells were imaged using a Leica TCS SP8 laser-scanning confocal microscope equipped with HyVolution deconvolution software. All images shown are either *en face* maximum projections through the full height of the apical domain, with the exceptions of X-Z sections, which are single plane confocal images. Live-cell imaging of the LLC-PK1-CL4 cells stably expressing GCaMP6 was performed with EVOS M5000.

### Image analysis

Image analysis was performed using ImageJ (NIH). For analysis of Myo7A microvillar targeting, *en face* images of polarized monolayers were inspected. Cells were binned as having targeting if microvillar tip-localized EGFP signal was observed in 3 or more microvilli on the apical surface. For determining the efficiency of targeting, we calculated the ratio of microvilli to cytosolic EGFP signal intensity as previously described (Crawley *et al*., 2014; Graves *et al*., 2020). Ratios at or below 1.5 were considered non-targeting, while above 1.5 were considered to have microvillar targeting. X–Z cross section images were used to quantify Myo7A microvilli to cytosolic signal intensity. For each construct, ∼30 cross section images were analyzed derived from ∼15 independent image Z-stacks. Microvilli:cytosol ratio values were generated as follows: for a given X–Z section, microvilli were first visualized using the F-actin (phalloidin) channel. The intensity of EGFP signal found in microvilli was measured at three to five microvilli in cells that were expressing the Myo7A construct. Another three to five intensity points were acquired from a region that corresponded to subapical cytosol. Intensity values from microvillar and cytosolic regions were averaged separately, and the resulting means were used to obtain microvilli:cytosol intensity ratios. For line scan analysis of hair cell targeting, a line was drawn parallel to the stereocilia axis across the rows using F-actin signal (visualized with phalloidin staining) as a reference and the intensity of Myo7A tagged with EGFP signal along that was recorded. Intensity values were normalized by the maximum gray scale value for an 8-bit image (i.e. 255). The corresponding positions for each intensity value was normalized so that the base of the stereocilia was equal to 0 and the tip was equal to 1. Normalized line-scans were then plotted together and fit using Prism v.6 software (GraphPad), which revealed the position of peak signal intensity and distribution width (S.D.) relative to the stereocilia. Data were analyzed in a blinded fashion.

### Calcium Ionophore (A23817) treatment of LLC-PK1-CL4 cells

LLC-PK1-CL4 cells stably expressing EGFP Myo7A WT full-length were seeded on coverslips at 80% confluency and allowed to grow for 4 days. Cells were then treated with A23817 (Sigma) at numerous concentration ranges and time courses. The concentration range of A23827 used was 1 μM, 4 μM, 7 μM, and 10 μM. Each concentration treatment of A23817 on CL4 cells was done for different time series (30 sec, 1 min, 5 mins, 10 mins, 1 hr, 4 hrs, and 10 hrs). Cells were then immediately fixed in 4% paraformaldehyde (Electron Microscopy Sciences) and processed as normal.

### Hair cell transfections

Experiments with mice were approved by the Institutional Animal Care and Use Committee of Indiana University -Indianapolis. Inner ear hair cell tissue explants were harvested from neonates at postnatal day P5 after decapitation and auditory hair cells were transfected with plasmid DNA encoding EGFP-tagged full-length Myo7A WT or R853C mutant by previously described injectoporation technique (Xiong *et al*., 2014). The tissue was dissected from postnatal day 5 C57BL/6 mice in Hank’s Balanced Salt Solution (HBSS, Life Technologies) and the cochlear duct was opened by making an incision between Reissner’s membrane and the stria vascularis. The tissue was adhered to to a plastic, tissue-culture treated dish (USA Scientific) containing DMEM/F12 (Thermo Fisher Scientific) with 1 mg/ml penicillin. For the injection step, a glass micropipette with a 2 mm tip diameter loaded with plasmid DNA (2 mg/ml in water) was inserted into the space between two IHCs, pressure was supplied by a microinjector to inject plasmid, and an ECM 830 electroporator was used to deliver a series of three 15 ms 60 V square-wave electrical pulses at 1 s intervals to platinum wire electrodes positioned over the injection site. After the electroporation, the culture media was exchanged with Neurobasal-A medium (ThermoFisher Scientific) supplemented with 2 mM L-glutamine (ThermoFisher Scientific), 1x N-2 supplement (ThermoFisher Scientific), 75 mg/ml D-glucose (ThermoFisher Scientific), and 1 mg/ml penicillin. Cultures were incubated for 18 hrs, then fixed with 4% formaldehyde in HBSS for 2 hrs and stained with Alexa Fluor 568 phalloidin (0.5 U/ml, Invitrogen) in PBS with 0.1% Triton X-100 (Sigma) at room temperature for 1 hr. The tectorial membrane was removed, and the tissue was mounted in Prolong Diamond (ThermoFisher Scientific). The cured slides were imaged with a Leica Plan Apo 63×/1.40 NA oil immersion objective on Leica SP8 inverted confocal microscope operating in resonant scanning mode (Leica Microsystems). Images were captured using Leica Application Suite X and deconvolved using Leica LIGHTNING deconvolution with the default settings.

### Statistical analysis

All graphs were generated and statistical analyses were performed using Prism v.6 (GraphPad). For all figures, error bars represent SD. Unpaired t tests were employed to determine statistical significance between reported values.

## Lead contact

Further information and requests for resources and reagents should be directed to and will be fulfilled by the lead contact Scott W. Crawley.

## Data availability

Data will be shared upon request to the lead contact.

---

**Figure S1.**
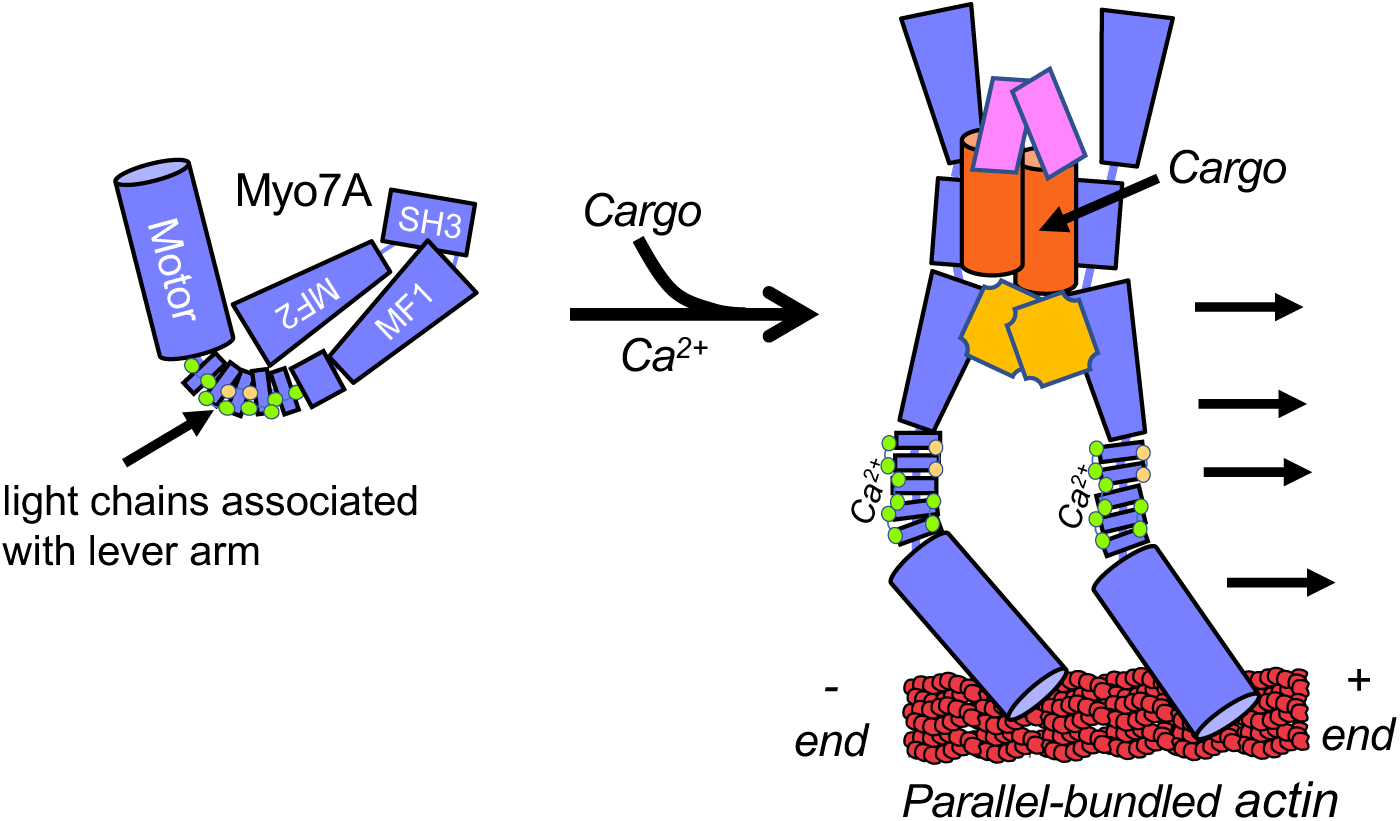
Current model of Myo7A regulation. Myo7A is thought to form a closed, autoinhibited conformation in which the tail folds back to inhibit the motor-neck region of the myosin. It has been previously proposed that cargo binding to the tail or Ca^2+^ binding to the light chains associated with the lever arm could activate the myosin. Cargo is also thought to oligomerize Myo7A.

**Figure S2.**
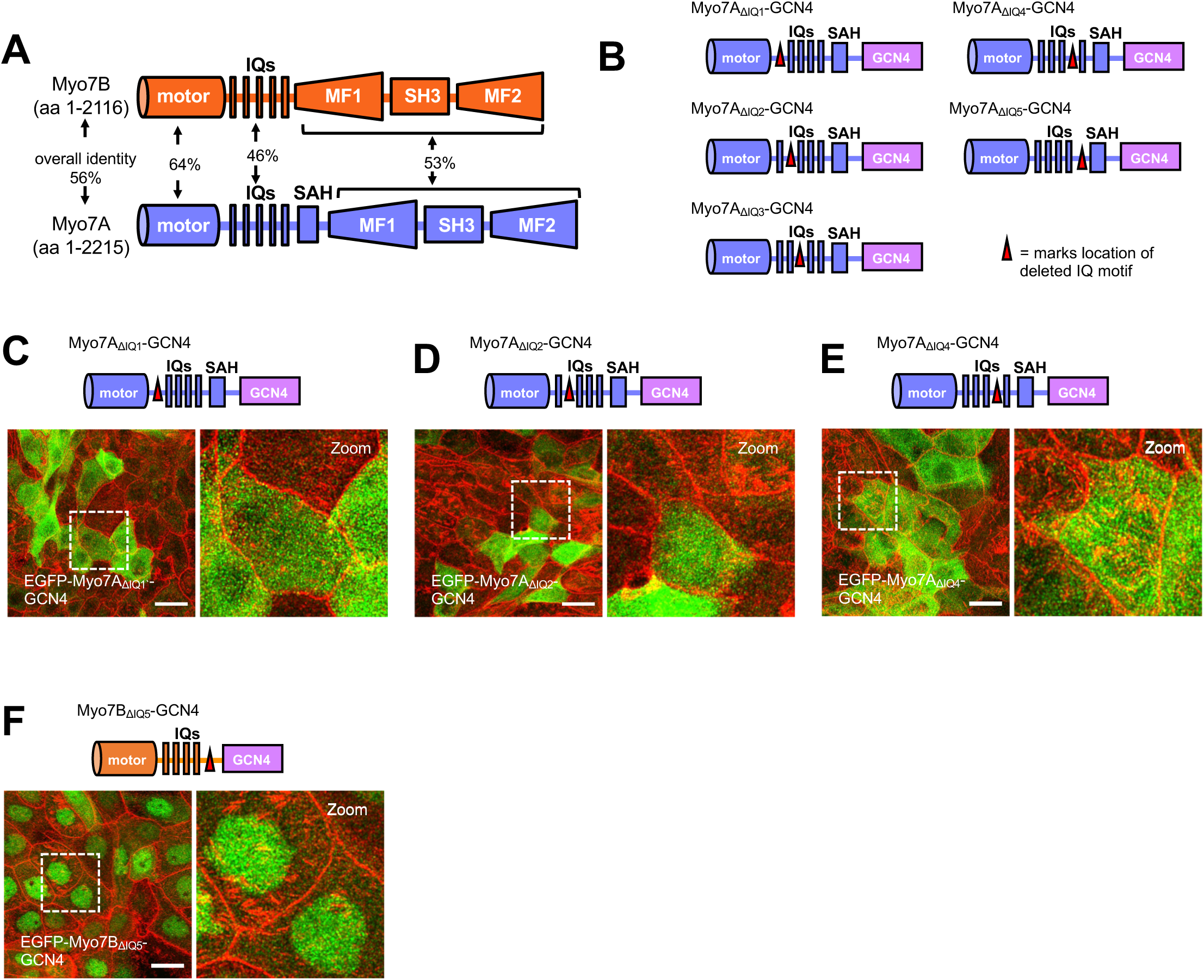
Myo7B does not exhibit the same lever arm regulation as Myo7A. (A) Diagram outlining the percent sequence identity shared between Myo7A and Myo7B. The lever arms exhibit the lowest identity between these homologous myosins. (B) Cartoon diagrams of the Myo7A forced-dimer constructs that contain internal IQ motif deletions used to map which IQ motif(s) could be controlling Myo7A targeting in epithelial cells. (C-E) Confocal images of CL4 cells stably expressing EGFP-tagged Myo7A forced-dimer containing an internal deletion of IQ1, IQ2 or IQ4. Cells were visualized for F-actin (red) and EGFP signal (green). Scale bars, 10 µm. (F) Confocal image of CL4 cell stably expressing EGFP-tagged Myo7B forced-dimer containing an internal deletion of IQ5. Cells were visualized for F-actin (red) and EGFP signal (green). Scale bars, 10 µm.

**Figure S3.**
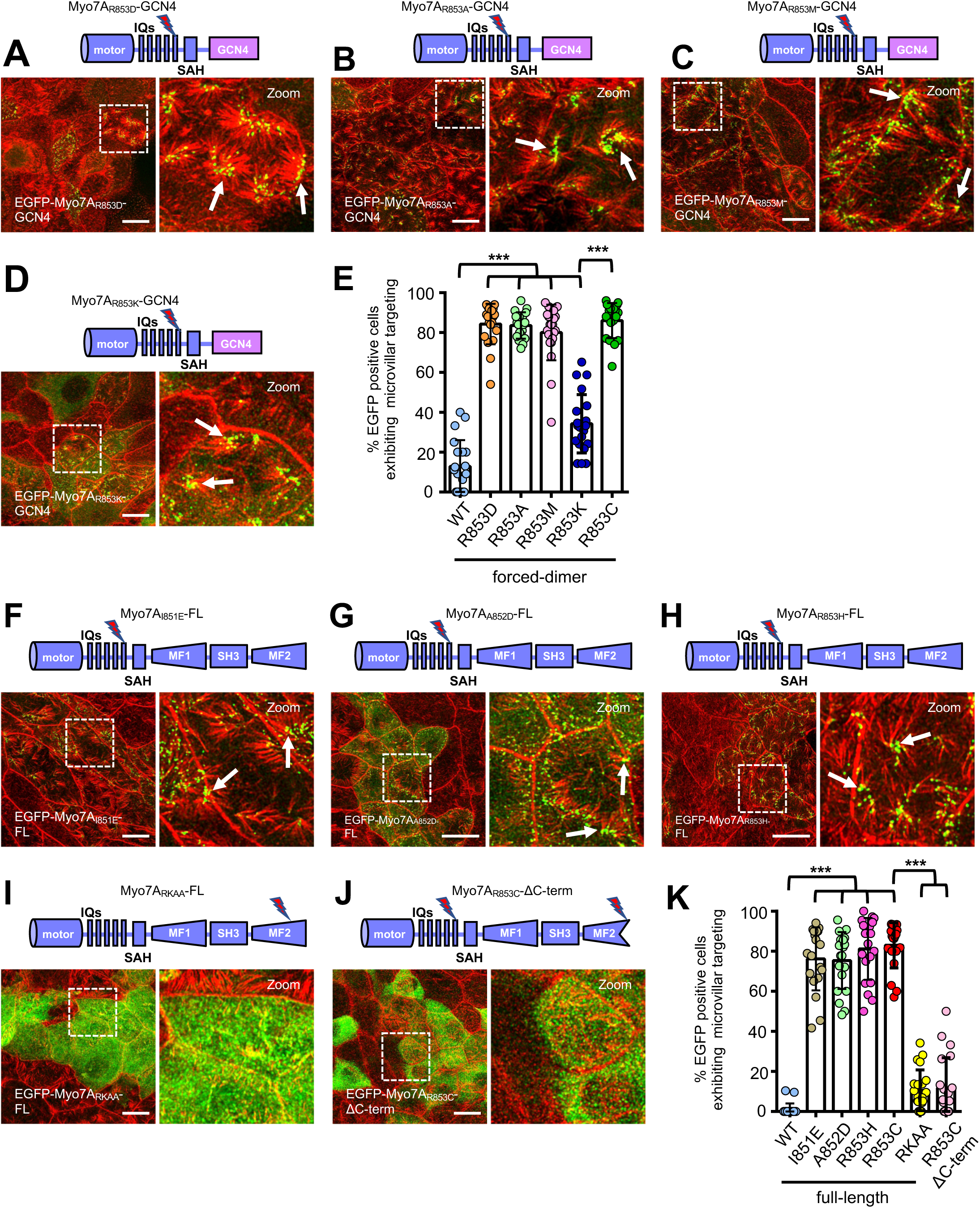
Activation of Myo7A in CL4 cells depends upon the specific mutation incorporated into IQ5 and a functional MF2 cargo binding domain. (A-D) Confocal images of CL4 cells stably expressing EGFP-tagged Myo7A forced-dimer that have different amino acids substitutions at the R853 position. Cells were visualized for F-actin (red) and EGFP signal (green). Scale bars, 10 µm. (E) Scatterplot quantification of the percent EGFP-positive cells exhibiting microvillar targeting of the mutant Myo7A forced-dimer constructs tested. ***p < 0.0001, t-test. Bars indicate mean ± SD. (F-J) Confocal images of CL4 cells stably expressing EGFP-tagged Myo7A full-length mutants constructs and a construct containing the R853C activation mutation along with a 33 amino acid truncation of the C-terminal MF2 domain (ΔC-term). Cells were visualized for F-actin (red) and EGFP signal (green). Scale bars, 10 µm. (K) Scatterplot quantification of the percent EGFP-positive cells exhibiting microvillar targeting of the mutant Myo7A full-length constructs and Myo7A R853C ΔC-term construct tested. Bars indicate mean and SD. ***p < 0.0001, t-test. Bars indicate mean ± SD.

**Figure S4.**
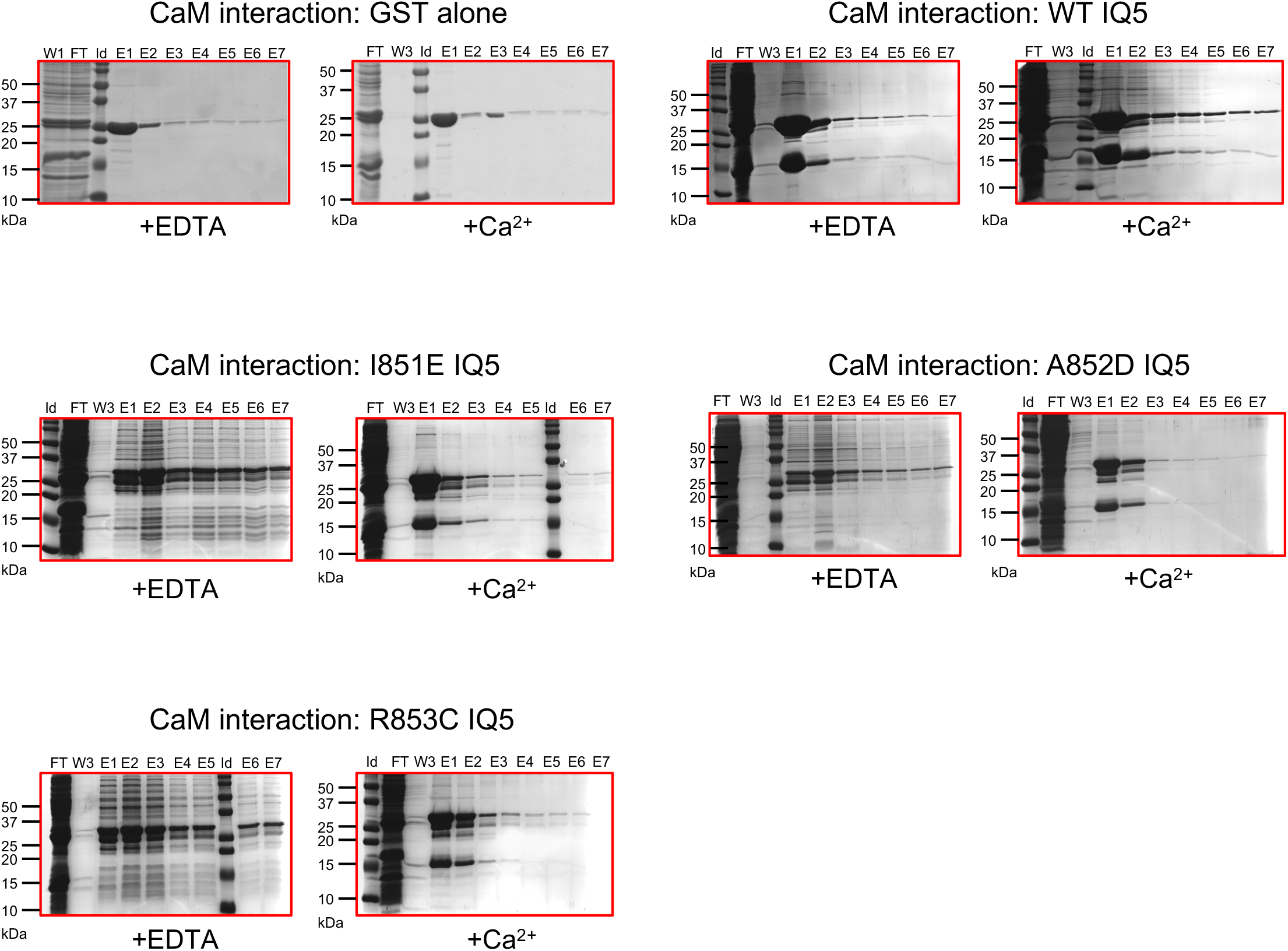
Uncropped Coomassie-blue stained SDS-PAGE gels showing the pull-down results between IQ5-activating mutants and CaM done in the presence and absence of Ca^2+^. Identity of the mutants are indicated above the gel sets. Also shown is the negative control testing the interaction of CaM with GST alone.

**Figure S5.**
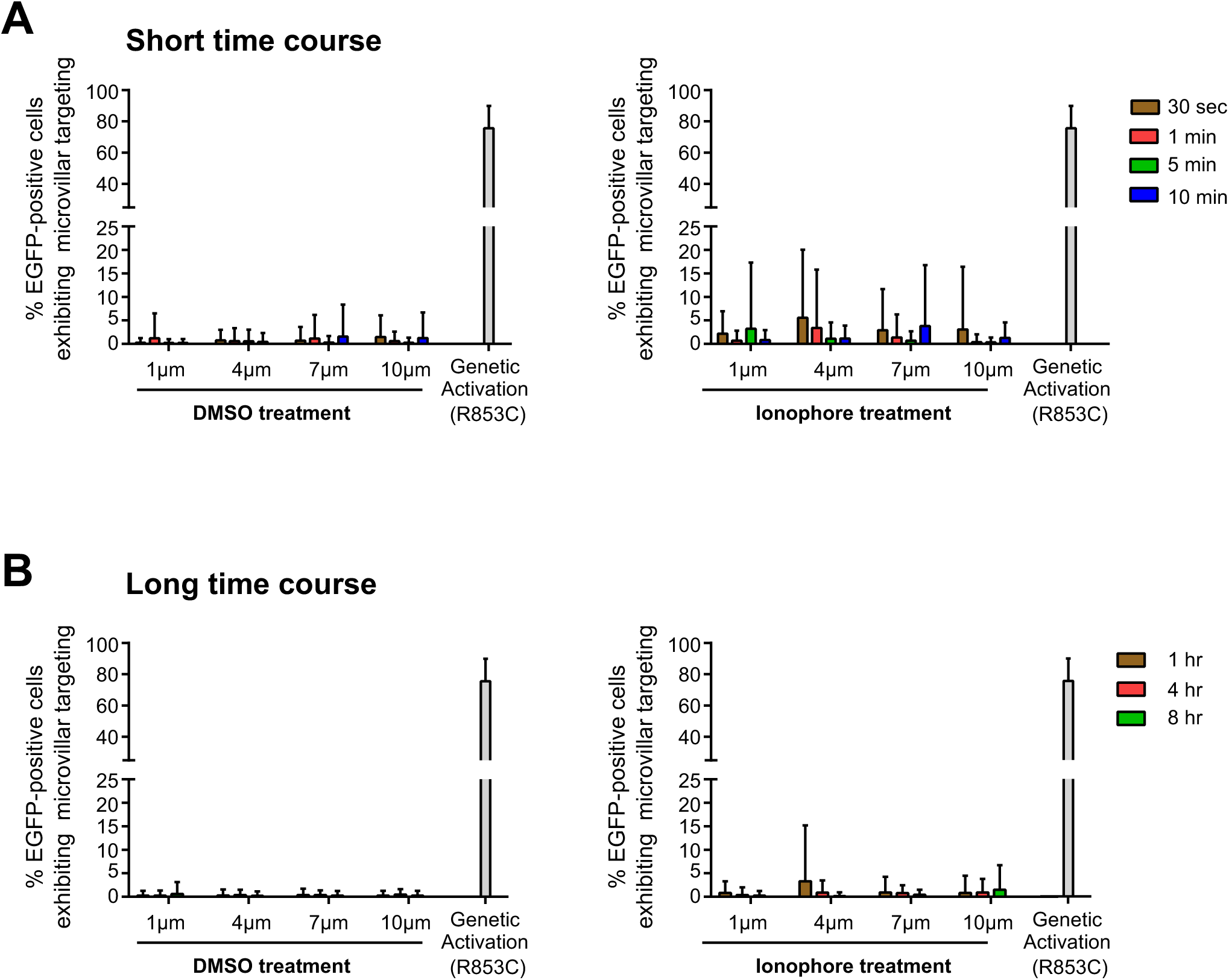
Exposure to Ca^2+^ ionophore A23187 does not potently activate Myo7A microvillar targeting in CL4 cells. CL4 cells stably expressing EGFP-tagged Myo7A WT full-length protein were exposed to different concentrations of Ca^2+^ ionophore A23187 and observed across both (A) short and (B) long time-courses. Microvillar targeting of the EGFP-tagged Myo7A R853C full-length construct was used as a comparison.

**Figure S6.**
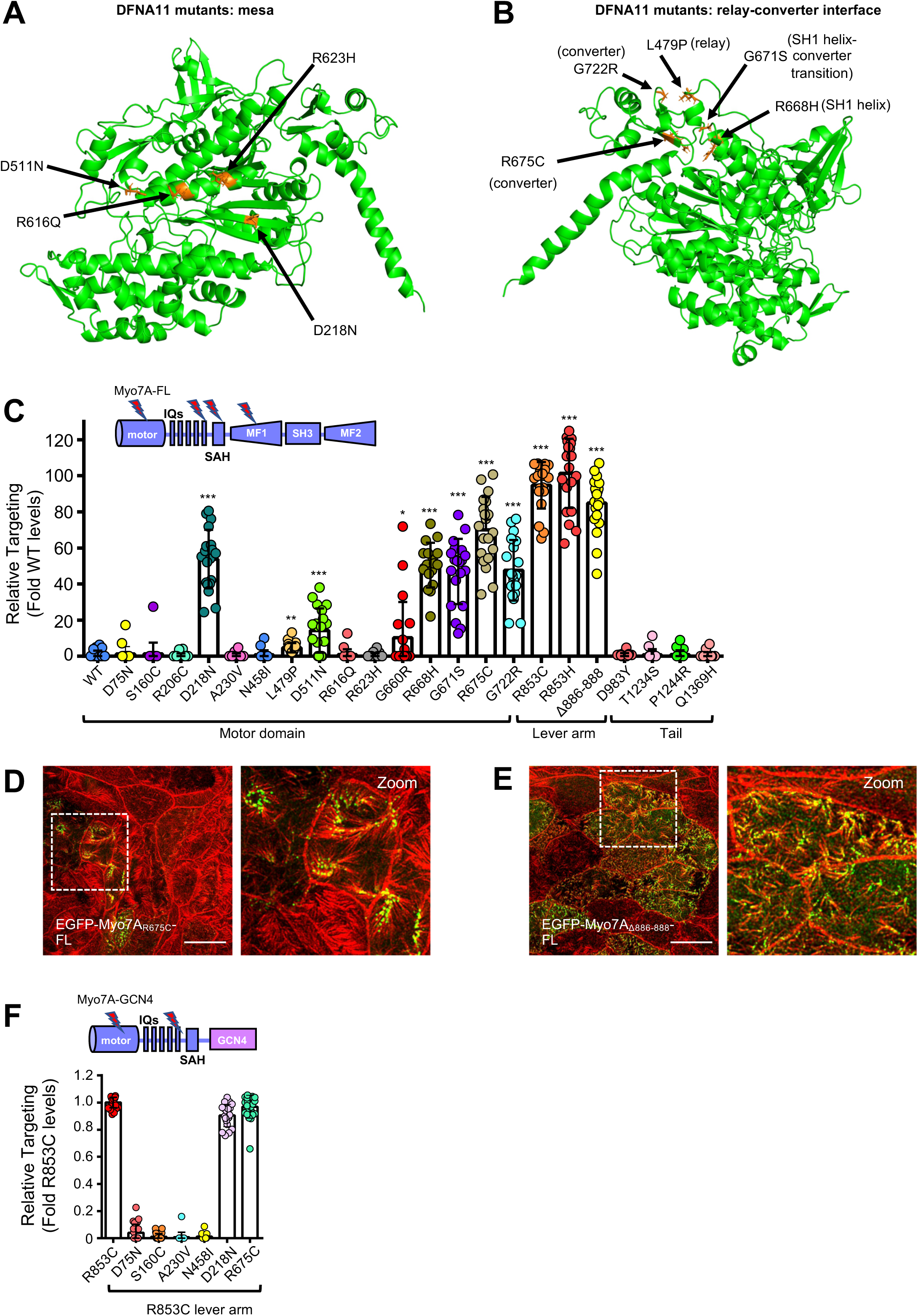
DFNA11 mutations that activate Myo7A map to distinct structural regions of the myosin. Structural models of Myo7A showing the location of DFNA11 mutants in the (A) myosin mesa and the (B) relay-converter domain interface that are able to activate Myo7A targeting in CL4 cells. The residues impacted by the DFNA11 mutations are labelled and shown in orange stick form. (C) Scatterplot quantification of the relative microvillar targeting of the DFNA11 variants compared to WT full-length Myo7A. All mutants were tested in the context of the Myo7A full-length construct. Bars indicate mean and SD. ns= not significant, *p < 0.01, **p < 0.001, ***p < 0.0001, t-test. (D-E) Examples of DFNA11 variants that activate microvillar targeting of the full-length Myo7A in CL4 cells. Cells were visualized for F-actin (red) and EGFP signal (green). Scale bars, 10 µm. (F) Scatterplot quantification of the relative microvillar targeting of the DFNA11 double mutant forced-dimer variants compared to Myo7A R853C forced-dimer alone. Bars indicate mean and SD. ns= not significant, ***p < 0.0001, t-test. Bars indicate mean ± SD.

**Figure S7.**
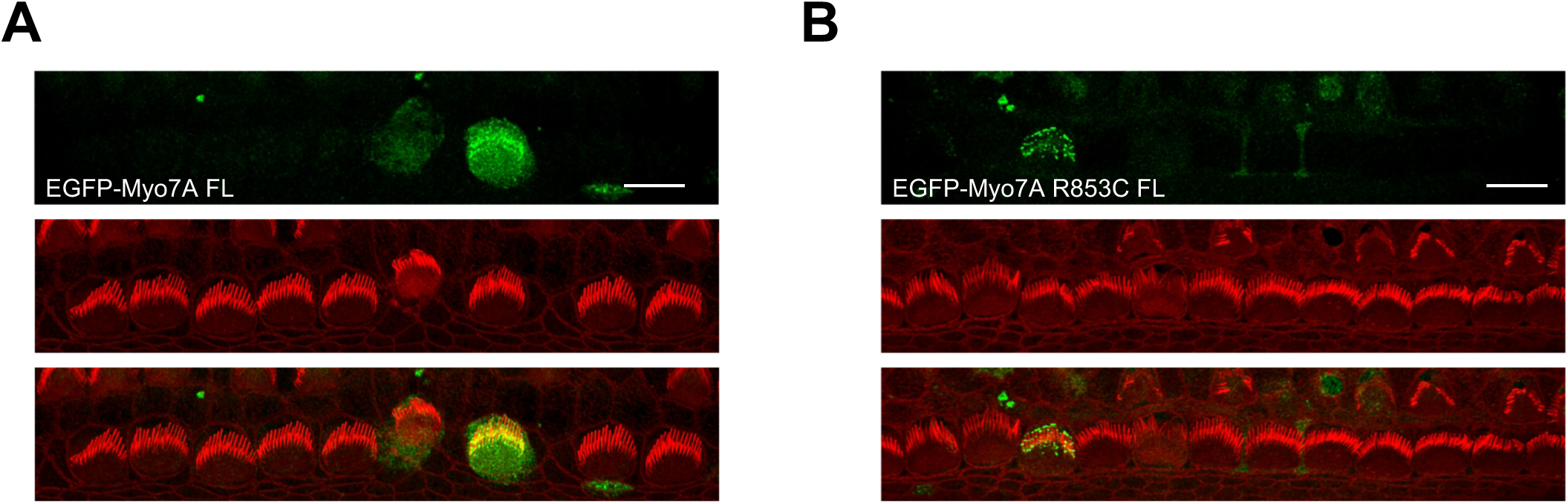
Additional examples of confocal images of (A) WT and (B) R853C Myo7A full-length transfections of mouse tissue explant cultured inner ear hair cells.

## Supplemental information

Document S1. Figures S1–S7 and Table S1

Video S1. Calcium Ionophore treatment of GCaMP6-expressing CL4 cells, related to Figure 5

---

**Table S1.**
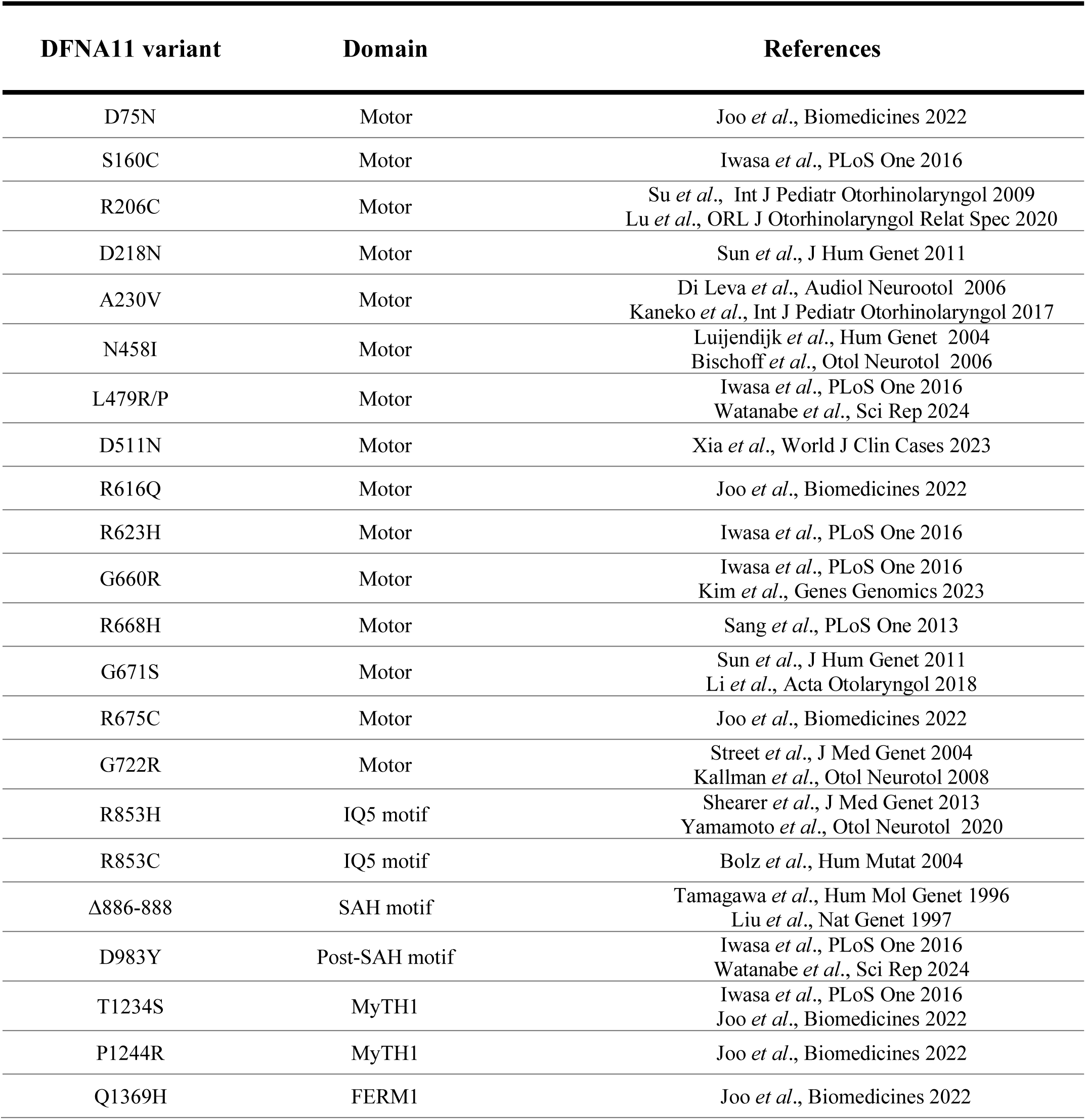

